# Echidna: A Bayesian framework for quantifying gene dosage effect impacting phenotypic plasticity

**DOI:** 10.1101/2024.12.15.628568

**Authors:** Joy Linyue Fan, Mingxuan Zhang, William O’Brien, Joshua D. Myers, Johannes C. Melms, Jana Biermann, Edridge D’Souza, Somnath Tagore, Nicolas Beltran-Velez, Kevin Hoffer-Hawlik, Alexander Preau, Isha Arora, Sharanya Chatterjee, Benjamin Izar, Elham Azizi

## Abstract

Phenotypic plasticity, the ability of cells to adapt their behavior in response to genetic or environmental changes, is a fundamental biological process that drives cellular diversity in both normal and pathological contexts, including in tumor evolution. While chromosomal instability and somatic copy number alterations (CNAs) are known to influence cellular states, it remains difficult to separate genetic from cell non-autonomous mechanisms that govern transcriptional variability. Here, we present *Echidna*, a Bayesian hierarchical model that integrates single-cell RNA sequencing (scRNA-seq) and bulk whole-genome sequencing (WGS) data to quantify the impact of CNAs on gene expression dynamics. By jointly inferring clone-specific CNA profiles and uncovering clonal dependencies, Echidna bridges genomic and transcriptomic landscapes within and across multiple time points, enabling the decoupling of gene dosage effects from cell-extrinsic factors on phenotypic plasticity. Applying Echidna to patient tumor specimens, we demonstrate its superior performance in clonal reconstruction and derive insights into resistance mechanisms.

## 1 Introduction

The interplay between the genome and transcriptome governs cellular behavior in both health and disease, driving complex biological processes such as differentiation, adaptation, and response to environmental changes [1–4]. An instructive example of this interplay is tumor evolution, where genomic instability and transcriptional variability lead to diverse phenotypic states, enabling cancer cells to adapt to selective pressures such as immune surveillance and therapeutic interventions [5–9]. However, the genome and transcriptome have largely been explored as disjoint features only.

With traditional genomics analyses, studies of tumor evolution using whole genome sequencing (WGS) have established that chromosomal instability and somatic copy number alterations play pivotal roles in the development and progression of cancer [10–13]. A well-understood mechanism by which this occurs is through the sequential accumulation of genetic alterations in genes such as tumor suppressors and oncogenes [14–18]. In this model, an initial set of mutations or CNAs in a single founder cell confers a selective advantage over healthy neighboring cells, leading to neoplastic proliferation. Over time, additional genetic alterations are acquired, eventually producing a lineage of genetically distinct clones with differential capabilities to survive and expand [19]. While these clones comprise one primary tumor, they may differ in their capacity to expand, metastasize, or resist treatments [20].

On the other hand, analyses of single-cell RNA sequencing (scRNA-seq) have focused on transcriptional variability and revealed extensive heterogeneity across phenotypic states, particularly within the tumor microenvironment (TME) [21–23], that are associated with disease progression or response to therapies [20, 24]. This transcriptional diversity reflects a phenomenon known as phenotypic plasticity, wherein a clone transitions between states to enhance survival (**Fig. 1a**). In the context of cancer, cellular plasticity enables malignant clones to evade immune response and therapy, leading to tumor proliferation [25]. The mechanisms that facilitate broader cellular plasticity are diverse, and include genetic, epigenetic, and transcriptional factors. For instance, mutations in key oncogenes, such as B-RAF, are required for transformation and tumor maintenance [26], yet epigenetic or transcriptional variability in the background of a fixed oncogenic mutation may drive critical phenotypes, such as resistance to BRAF-directed targeted therapies [27]. It is unclear to what extent such plasticity at the transcriptional level is enabled by gene dosage effects determined by CNAs in the driver genes or other genomic loci. Given that CNAs, chromosomal instability (CIN), and aneuploidy are hallmarks of solid cancers [28, 29], quantitative tools to determine whether transcriptional variability is primarily driven by intrinsic genetic features (gene dosage effect [30]) or external stimuli and interactions with the microenvironment are critically important.

**Figure 1:**
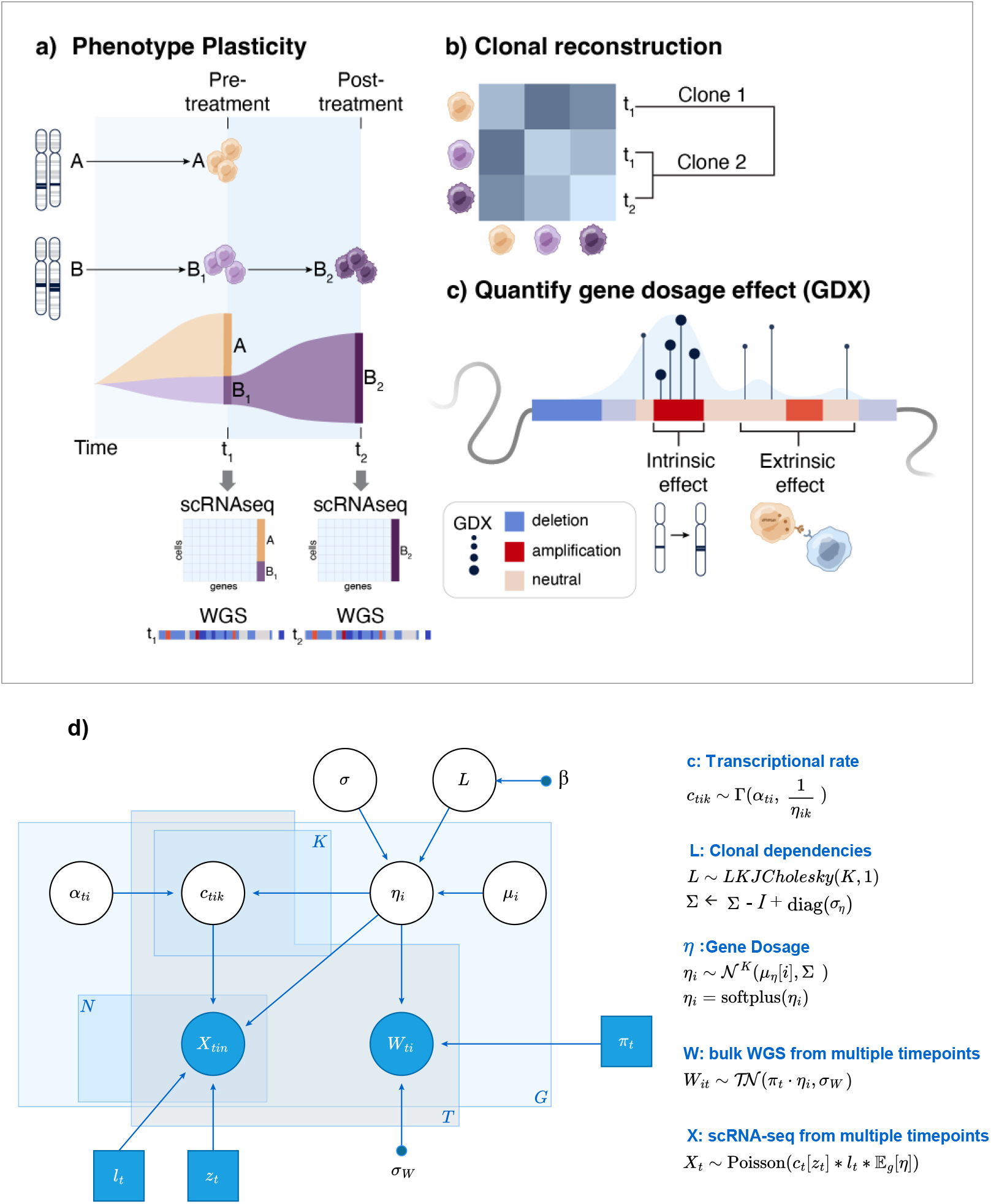
The Echidna model. (a) Echidna integrates single or multiple time point scRNA-seq and WGS data in order to characterize the driving forces of phenotypic plasticity and clonal expansion. (b) Echidna infers cluster-level gene dosage values, while also learning the correlation between clusters to reconstruct the underlying clonal structure. (c) Echidna’s hierarchical structure allows for the computation of a novel metric, the GDX, that quantifies the extent to which a phenotypic shift in gene expression (shown in bars) may be attributed to a genetic alteration. The outputs of Echidna may be used to identify hotspots of differentially-expressed (DE) genes, while identifying their dependence on genetic alterations through GDX (shown with dot size). (d) Plate model of Echidna. White nodes represent latent random variables and shaded nodes represent observed random variables. Shaded squares represent parameters we fix according to observed data. Arrows indicate conditional dependency relationships and the direction represents the generative process. Plates indicate dimensions of independence.

This gap motivates the development of computational tools that disentangle the impact of intrinsic genetic features from extrinsic cues, such as cellular interactions and their spatial localization, which dynamically shape cellular states. Previous attempts to quantify the link between the genome and transcriptome [31, 32] relied on bulk sequencing, which obscures tumor heterogeneity and the contribution of non-cancerous cells in the microenvironment. While paired genomic and transcriptional profiling at the single-cell level is technically possible [33–36], such data are difficult to generate at scale and depth, and datasets coupling these methods remain scarce [37, 38]. Conversely, scRNA-seq analysis pipelines for clustering single-cell transcriptomics and identifying differentially-expressed genes are also limited in the scope of biological insights as they merely focus on phenotypic plasticity and do not elucidate driving factors such as gene dosage, clonal identities, and lineages. For example, in the cancer context, numerous studies have defined transcriptional programs associated with response or resistance to therapies [39, 40]; however, it is unclear if these programs are up-or down-regulated because of specific genetic aberrations (e.g. amplification or deletion) or extrinsic factors. This distinction is key for generalizability to larger clinical cohorts and guiding therapeutic solutions.

Methods for inferring single-cell CNAs from scRNA-seq data ([41],[42],[43]) have demonstrated the success of using sequential assumptions in a Hidden Markov Model (HMM) framework on gene expression data to infer latent copy number states along the chromosome. However, this sequential constraint also restricts the ability to additionally model the relationship between CNAs and expression, as the two do not have the same sequential dependencies. Thus, these methods focus primarily on CNA and phylogeny inference, without considering downstream transcriptomic effects. Additionally, these methods require normal cells as a reference for the reconstruction of CNAs, which may introduce biases if the selected population of cells is not suitable. Finally, previous approaches also fail to model temporal dependencies in datasets collected across multiple time points. The temporal dimension is critical for revealing the impact of CNAs on clonal evolution over time, as well as unveiling the genomic impact on phenotypic plasticity.

To address these challenges, we present *Echidna* (Examination of Clone plasticity using Hierarchical modelIng and Deconvolution of copy Number Alterations), a Bayesian hierarchical model designed to bridge the gap between the genome and transcriptome using single or multiple time-point datasets, and quantify the dependence between transcription and CNAs. Echidna requires only scRNA-seq and bulk WGS, both of which are readily obtainable, including from routine and archived clinical specimens. We demonstrate that by integrating expression data with WGS, Echidna not only enables improved inference of clone-specific CNAs and clonal dependencies without the need for a normal reference or underlying sequential assumptions, but the generative hierarchical framework also captures the complex underlying biological processes linking tumor evolution to plasticity. We anticipate that insights from Echidna will enable both the discovery of novel genome alterations driving tumor plasticity and resistance to therapy, as well as the transcriptional mechanisms through which these alterations take effect.

## 2 Results

### 2.1 Overview of Echidna

Echidna is a Bayesian hierarchical model designed to study the dynamic impact of genotype on transcriptional variability, phenotypic plasticity, and clonal evolution. Specifically, Echidna provides a robust framework for quantifying the influence of genetic features defined by CNAs on phenotypic plasticity. Echidna integrates two paired data modalities, WGS and scRNA-seq data, from samples collected at single or multiple time points.

Echidna takes in identified phenotypic states as *clusters* of cells using standard clustering approaches applied to scRNA-seq data as input. When longitudinal data is available from the same biological system or patient, single-cell transcriptomes from all time points are merged prior to clustering to allow for downstream detection of evolving clones. The input scRNA count tensor across time points can be represented as *X*^*T ×N×G*^ for *T* time points, *N* cells, and *G* genes. We also denote the observed proportion of clusters within a sample or time point as *π ∈* ℝ^*T ×K*^, with *K* being the total number of clusters. Finally, WGS data providing CNA estimates at the level of genomic regions is mapped to individual genes to obtain bulk-level *gene dosage*, denoted as *W* ^*T ×G×*1^ across *G* genes and *T* time points.

The generative process built into the Echidna model consists of two main components (**Fig. 1a**). First, we assume bulk gene dosage measurements from WGS are a weighted sum of unknown (latent) cluster-specific gene dosage profiles (*η*), with the weights corresponding to the proportion of clusters (*π*) in scRNA-seq data in each time point (*t*). To enable the inference of clonal structure and evolution, we allow correlations between cluster-specific gene dosage profiles (**Fig. 1b**). Specifically, gene dosage profiles are modeled as multivariate Gaussian distributions centered around neutral copy number (2) with a learned covariance random variable Σ that represents cluster correlations (see **Methods**).

Second, single cell transcript count is modeled as a Gamma-Poisson distribution to account for overdispersion, similar to other probabilistic models for scRNA-seq data [44, 45]. Importantly, to link the two components and quantify the influence of gene dosage on gene expression in each cluster, we define *transcriptional rate* as a latent parameter (*c*). This parameter follows a Gamma distribution with a rate dependent on gene dosage. Finally, the Poisson rate of expression is constructed to be a function of the latent transcriptional rate as well as cluster and library size biases (see **Methods**).

Integrating transcriptomic and WGS data improves accuracy in learning CNAs compared to methods that use scRNA data alone, as the bulk gene dosage measures set a constraint on the sum of CNAs across cells. With a hierarchical structure, Echidna jointly learns cluster dependencies that inform cluster-specific gene dosage and, in turn, its impact on expression. Cluster dependencies are therefore informative of clonal structure (**Fig. 1b**). After fitting the model presented in **Fig. 1d**, the following post-processing steps are performed to obtain CNAs and clonal architecture: We incorporate genomic position information by first smoothing the learned gene dosage matrix (*η*) with a 1D Gaussian kernel along the genome. To derive cluster-specific neutral values, a Gaussian Mixture Model (GMM) is fitted to smoothed copy number values inferred for each cluster (**Fig. S1a**). The gene dosage values are then adjusted by the neutral value, defined as the mode of the most frequent GMM component in the cluster (see **Methods**). Lastly, a Gaussian HMM model is applied to predict copy number states in genomic regions (**Fig. 1c, S1b**). Hierarchical clustering can then be performed on either the inferred correlation between gene dosage values of clusters or the estimated posterior covariance directly, depending on sequencing depth and noise in the data (see **Methods**). From the hierarchical clustering, we obtain a proxy for a phylogenetic tree between clusters. The tree is then used to define *clones* as groups of clusters with similar CNAs. We perform threshold-based cutting on the tree to obtain the flat cluster labels corresponding to clones.

A key feature of Echidna is its ability to track clones and their phenotypic plasticity over time. To achieve this interpretability, while avoiding full covariance matrix inference, we parameterize the covariance random variable Σ as a combination of off-diagonal elements given by its lower Cholesky factorization (*L*) and diagonal elements given by cluster-specific variance (*σ*). The off-diagonal structure is sampled from an *LKJ* distribution [46], ensuring robust and numerically stable inference of dependencies between clusters from the same or different time points. The *LKJ* prior also allows tuning the degree of correlation between clusters through its shape hyperparameter (*β*), adjusting the resolution of subclonal structures (see **Methods**). As a result, clones that exhibit different phenotypic states within the same or across multiple time points can be identified as clusters with strong similarities in CNAs and are subsequently merged into a single clone.

Finally, to study the influence of gene dosage on expression, we define a metric for *gene dosage effect* (GDX). Specifically, we use posterior distributions learned by Echidna to quantify the dependence of transcriptional variability (*c*) on gene dosage (*η*) in the form of explained variance between hierarchically-linked random variables (see **Methods**). This metric can be computed for any individual gene or geneset to assess if their differential expression is strongly dependent on gene dosage or conversely likely to be driven by extrinsic effects (**Fig. 1c**). We further pinpoint gene signatures with significantly stronger links between expression and gene dosage in subpopulations of cells using a rank-based permutation test. A set of genes are first ranked by GDX. Enrichment scores are then computed similar to GSEA [47, 48]. To assess the statistical significance of enrichment scores, a permutation test is performed to obtain p-values (see **Methods**).

### 2.2 Echidna outperforms other methods in clonal reconstruction

To assess Echidna’s ability to infer clonal architecture, we applied it to matched single-nucleus (sn) hybrid DNA and RNA-seq datasets with corresponding WGS collected from frozen tissues from endometrial cancer patients [49] (**Fig. 2a**). We limit our analyses to patients 2 and 5 described in [49], as these tumor specimens involved significant genetic differences between clones as well as phenotypic diversity [49]. Sequenced using a novel adaptation of the BAG technology [49], the paired DNA-RNA measurements allowed performing ground truth validation of our results. Specifically, we fit our model to the WGS and snRNA-seq, and reserved the snDNA-seq data for validation.

**Figure 2:**
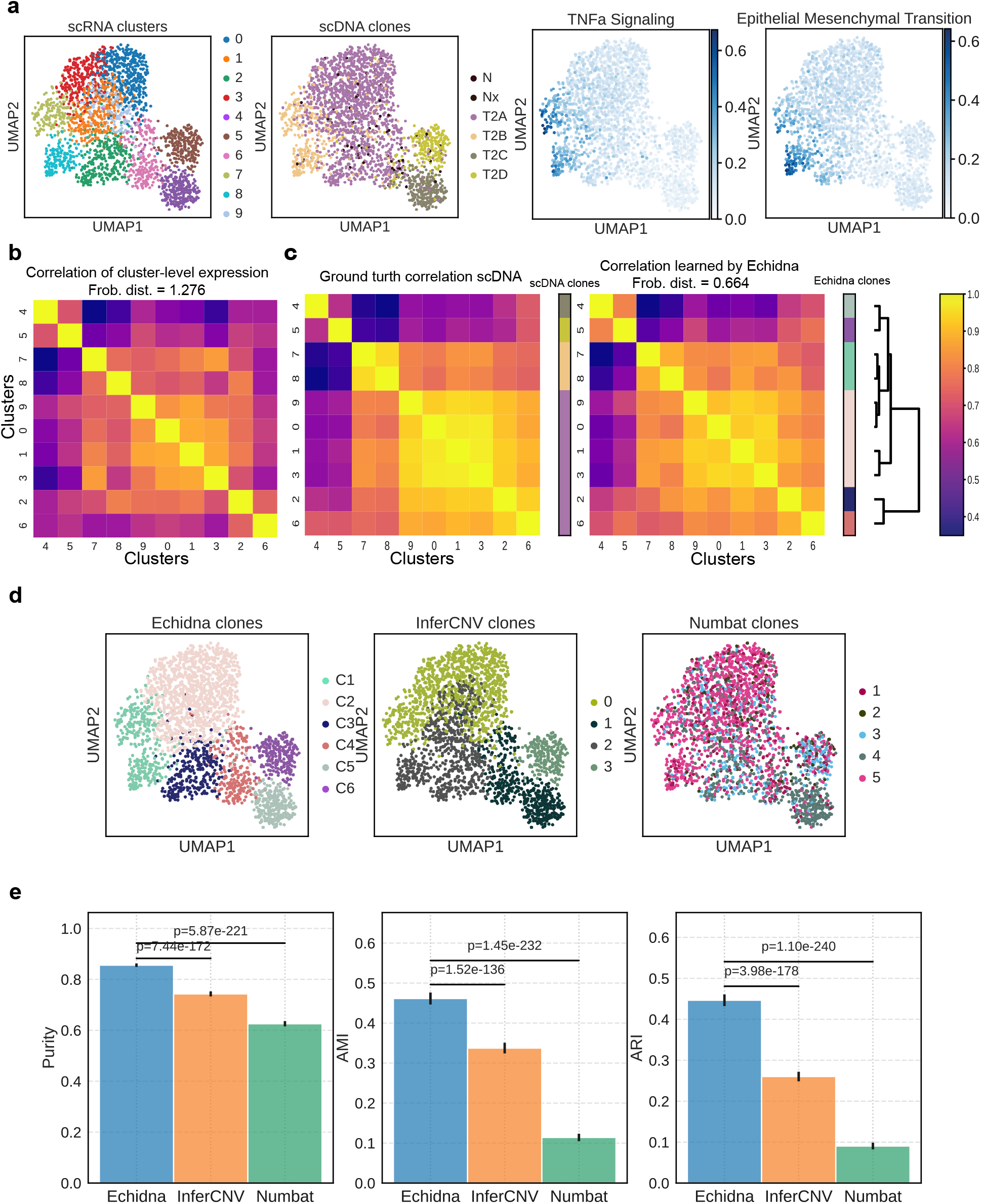
Echidna achieves accurate clone reconstruction by learning cluster-specific gene dosage and cluster correlation structure. **(a)** UMAP of the scRNA-seq data from endometrial cancer patient specimens [49] where each dot represents a cell. The cells are colored by Leiden clusters identified from expression (left), clones identified by matching scDNA data (middle), and expression of gene signatures enriched in clone T2B exhibiting phenotypic plasticity (right). **(b)** Correlation matrix of pseudo-bulked scRNA expression at the cluster level. Frobenius distance to the ground truth scDNA correlation is computed. **(c) Left**: The empirical correlation matrix of scDNA data across clusters as ground truth clonal dependencies. Colorbar shows the reported grouping of clusters into ground truth clone structure [49]. **Right**: The learned correlation matrix from Echidna. Colorbar shows clones identified by Echidna through hierarchical clustering and tree cutting. Dendrogram represents the learned tree grouping cell clusters. **(d)** UMAP of the scRNA-seq data colored by clones identified by Echidna, InferCNV, and Numbat respectively **(e)** Benchmarking clone reconstruction between Echidna, InferCNV, and Numbat. The metrics used are purity, adjusted mutual information (AMI), and adjusted rand index (ARI). Error bars computed from 100 bootstrapping iterations are on top of the bar graphs and p-values are computed with t-tests between methods.

**Figure S1:**
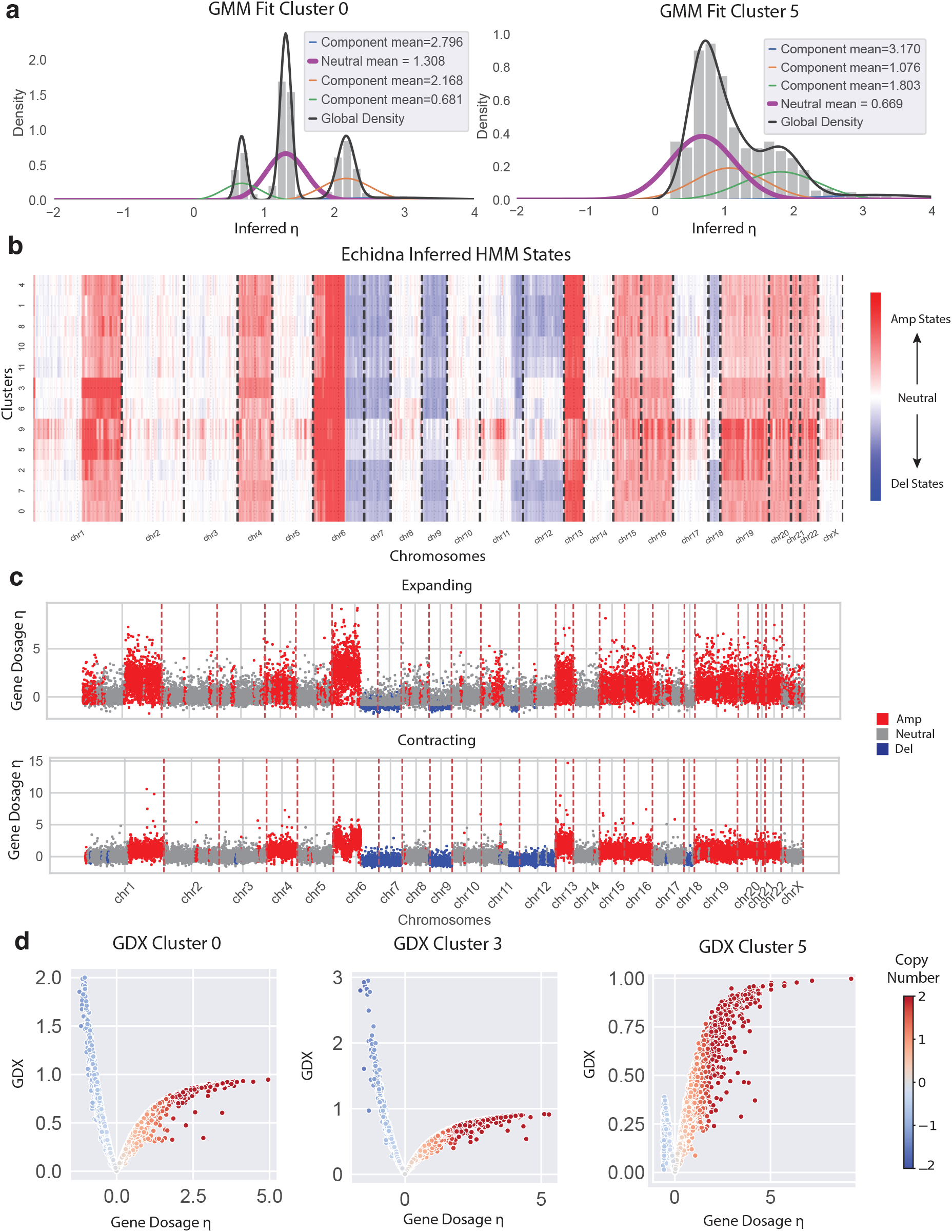
Echidna utilizes a Gaussian Mixture Model (GMM) and Gaussian HMM in post-processing steps. (a) Echidna fits a GMM to inferred copy number values per cluster. The GMM component with the most genes assigned (purple line) is used as the neutral mean. (b) HMM-inferred copy number states inferred from smoothed copy number values per cluster. Color represents different levels of amplified vs deleted states. (c) HMM-inferred copy number states based on learned gene dosage values (y=axis) in clusters grouped by temporal dynamics (expanding or contracting over time). Amp=amplification. Del=deletion. (d) Gene Dosage Effect (GDX) computed per cluster for a melanoma specimen as example [20]. The scatterplot demonstrates the relationship between GDX (y-axis) and gene dosage (x-axis and color). Each dot represents a gene and is colored by gene dosage.

**Figure S2:**
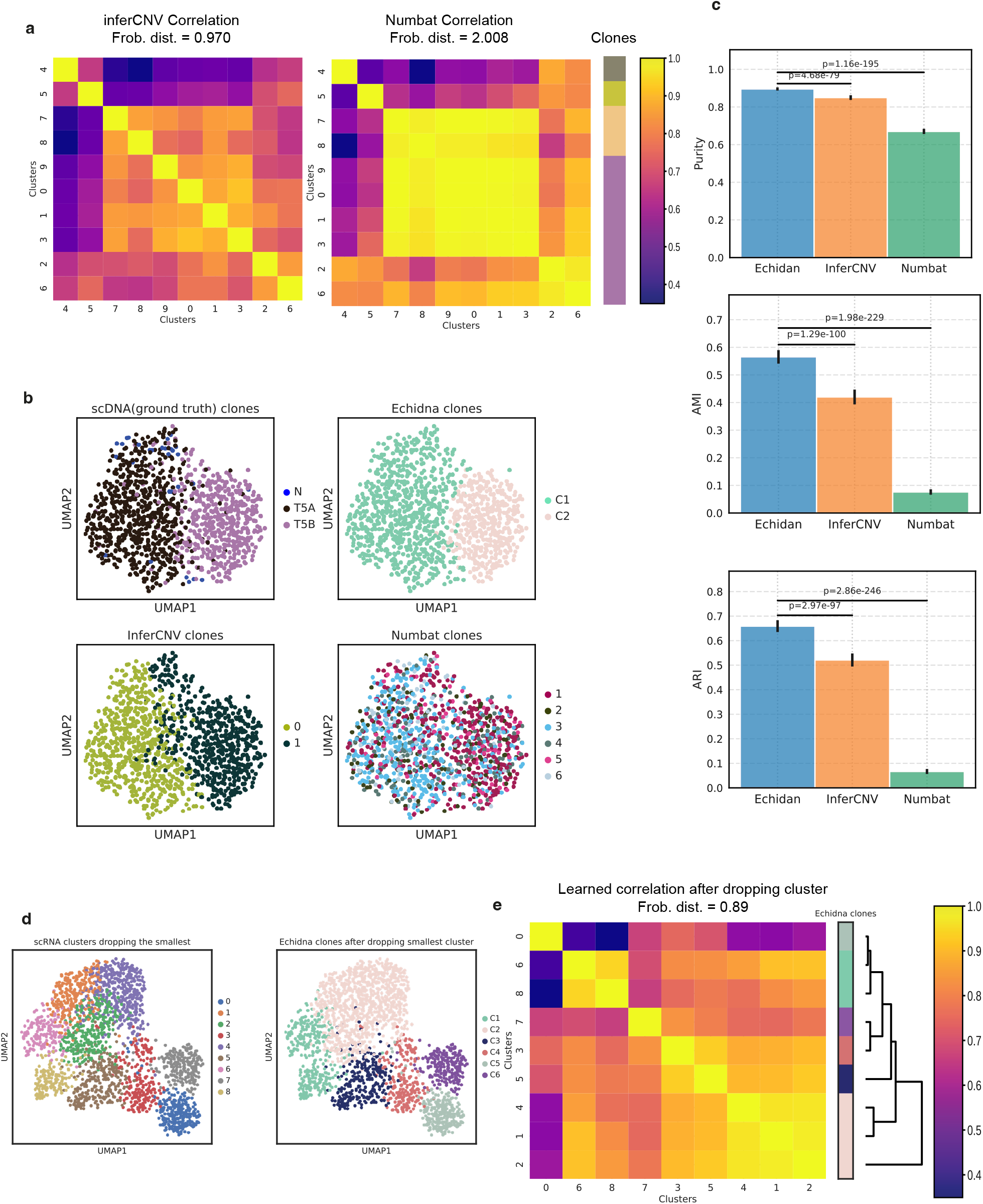
Echidna’s ability to reconstruct clones is generalizable and demonstrates robustness to noisy or small clusters. **(a)** Correlation between clusters recovered by InferCNV and Numbat for endometrial cancer tumor 2. Frobenius distances are computed to the ground truth correlation matrix from scDNA data. **(b)** UMAP of scRNA-seq data of an independent tumor 5. Each dot is a single cell. Colored by scDNA clones and clones detected by Echidna, InferCNV, and Numbat, respectively. **(c)** Benchmarking Echidna’s clone reconstruction with InferCNV and Numbat on tumor 5. The metrics used are purity, adjusted mutual information (AMI), and adjusted rand index (ARI). Bootstrapped error bar are at the top of the bar graphs and p-values are computed with t-tests between methods.**(d)** UMAP of scRNA-seq data of tumor 2 while dropping the smallest cluster to test method robustness. Cells are colored by Leiden cluster labels (left) and clones recovered by Echidna (right). **(e)** Correlation matrix inferred by Echidna on tumor 2 data with the smallest cluster dropped. Echidna clone calls are shown. Dendrogram shows the grouping of cell clusters.

Examining the clones identified by scDNA data in tumor 2, we observe that some clones with similar CNAs exhibit heterogeneous phenotypic states. For example, clone T2B corresponds to distinct clusters 7, 8 in scRNA-seq data that display functionally different transcriptional signatures from each other. Differential gene expression analysis reveals that the T2B clone is enriched in genes related to TNF-alpha signaling (*CCL20, TIPARP, NR4A3*) compared to other clones, while cluster 8 is enriched in genes involved in epithelial-mesenchymal transition (*COL7A1, DACT2, FSTL3, LAMA1*) compared to cluster 7 (**Fig. 2a**). Because of these transcriptional differences, computing the correlation between clusters using scRNA expression data alone does not reflect a strong similarity between clusters corresponding to the same clone (**Fig. 2b**). On the other hand, similarities between clusters at the CNA-level measured by scDNA (**Fig. 2b**) reveal a drastically different structure (quantified by Frobenius distance). We demonstrate that the integrative nature of Echidna’s deconvolution successfully recovers cluster-level genetic similarities that more closely resemble the ground truth scDNA clones while retaining some variability within clones (**Fig. 2c**). For example, clone T2A is further split into subclones (C2, C3, C4) to reflect its phenotypic plasticity.

To benchmark our results, we compared performance in reconstructing clones to commonly used state-of-the-art methods, namely InferCNV and Numbat. Using scDNA clones[49] as ground truth labels, we see that Echidna outperforms both methods in clonal reconstruction (**Fig. 2d**). For instance, we observe that while inferCNV captures general clonal architecture, it incorrectly groups cluster 8 with 2 into one clone (**Fig. 2d**), likely due to their transcriptional similarity (**Fig. 2b; Fig. S2a**), and does not detect its genetic similarity to cluster 7. As another example, we notice that Numbat captures a global clonal structure without identifying distinct clones. It groups clones T2A and T2B into the same cluster (**Fig. 2a,d**) and fails to detect the genetic difference between these clones (**Fig. 2d**).

In contrast, Echidna learns a covariance matrix that more accurately approximates the clonal structure compared to Numbat and InferCNV, as quantified by the Frobenius distance (**Fig. 2c; Fig. S2a**). This is partly due to Echidna’s integration with WGS data, which is readily obtainable and serves as a genetic reference in cases where the scRNA-seq may be of lower depth. As noted in [49], the coverage of the novel hybrid sequencing method is lower than standard single-nucleus sequencing methods for DNA and RNA separately.

For a more systematic evaluation, we quantify the quality of clone reconstruction by comparing clustering accuracy where the ground truth cluster labels are given by the scDNA clones (see **Methods**). To obtain Echidna clones, we assign labels based on thresholding the dendrogram of the correlation matrix between clusters. To obtain inferCNV clones, we performed K-means clustering with the single cell level CNA estimates similar to our previous work [20]. Numbat clones are retrieved directly from the single-cell clone assignment output. We computed three standard clustering metrics to compare performances: *purity* which measures the extent to which inferred clones contain only cells from ground truth clones; *AMI* (Adjusted Mutual Information) which quantifies the agreement between the inferred clones and ground truth labels while adjusting for chance; and *ARI* (Adjusted Rand Index) which measures similarity between inferred clones and ground truth by considering all pairs of cells and counting pairs that are assigned in the same or different clones, while adjusting for chance(see **Methods**). We observe significant improvement with Echidna compared to other methods with all metrics (**Fig. 2e**; *p <* 1*e* − 10). We then applied Echidna to a second tumor sample (from patient 5 in [49]) with a less complex clonal structure and performed systematic benchmarking with InferCNV and Numbat again to assess the generalizability of our model. We again observe improved performance with Echidna against both methods in terms of clone reconstruction (**Fig. S2b,c**; *p <* 1*e* − 10).

To demonstrate the robustness of our results, particularly to small or noisy clusters, we dropped the cluster with the fewest number of cells in tumor 2, and reproduced the ability to detect the clone with the greatest phenotypic plasticity (C1; **Fig. S2d**) and reconstructed cluster dependencies that resemble ground truth clonal structure (**Fig. S2e**). We do this by refitting Echidna on the scRNA-seq data with the smallest cluster removed using an identical set of hyperparameters and performing the same post-processing steps to obtain clone structures. Echidna successfully recovered the identical clone structure as observed prior to the single cluster removal (**Fig. S2d**,**e**).

### 2.3 Echdina achieves robust estimation of gene dosage and accurate prediction of CNV states

In addition to evaluating performance in clonal structure, we benchmarked Echidna’s ability to estimate CNA profiles at the level of phenotypic clusters against other methods. Our gene dosage estimates from Echidna revealed distinct clone-specific CNA events that match ground truth scDNA profiles (**Fig. 3a**). Specifically, we note amplifications in chromosome 1 in clusters 4, 5; amplifications in chromosome 4 in cluster 5; and deletions in chromosomes 7, 8, and 9 in clusters 7, 8 (which were previously identified to be from the same clone; **Fig. 2c**). To evaluate the quality of CNA estimates, we compared predicted cluster-level CNA estimates from each method to the scDNA ground truth. We utilize similar preprocessing of the scDNA data as [49], binning the data into 300 genomic bins to improve the signal. In order to benchmark the CNA estimates, we obtained cluster-level CNA estimations per gene from all methods and grouped them to match the genomic bins used by [49] in the scDNA data. For Echidna, we directly used the posterior mean of our gene dosage random variable. For InferCNV, we aggregated single-cell CNA estimates into pseudo-bulk profiles to generate CNA estimations at the cell cluster level. For Numbat, we constructed cluster-level CNA estimates from clone-level maximum a posterior (MAP) total copy number ratio relative to diploid. To measure reconstruction quality, we use Pearson correlation coefficient (*PearsonR*) between estimated CNA profiles and the scDNA ground truth. Echidna outperforms both InferCNV and Numbat in terms of CNA profile estimation and was able to detect clone-specific patterns (**Fig. 3a,b; Fig. S3a**). We conducted similar benchmarking experiments on an independent sample (tumor 5) and observed that Echidna outperformed other methods on this dataset as well(**Fig. S3b**).

**Figure 3:**
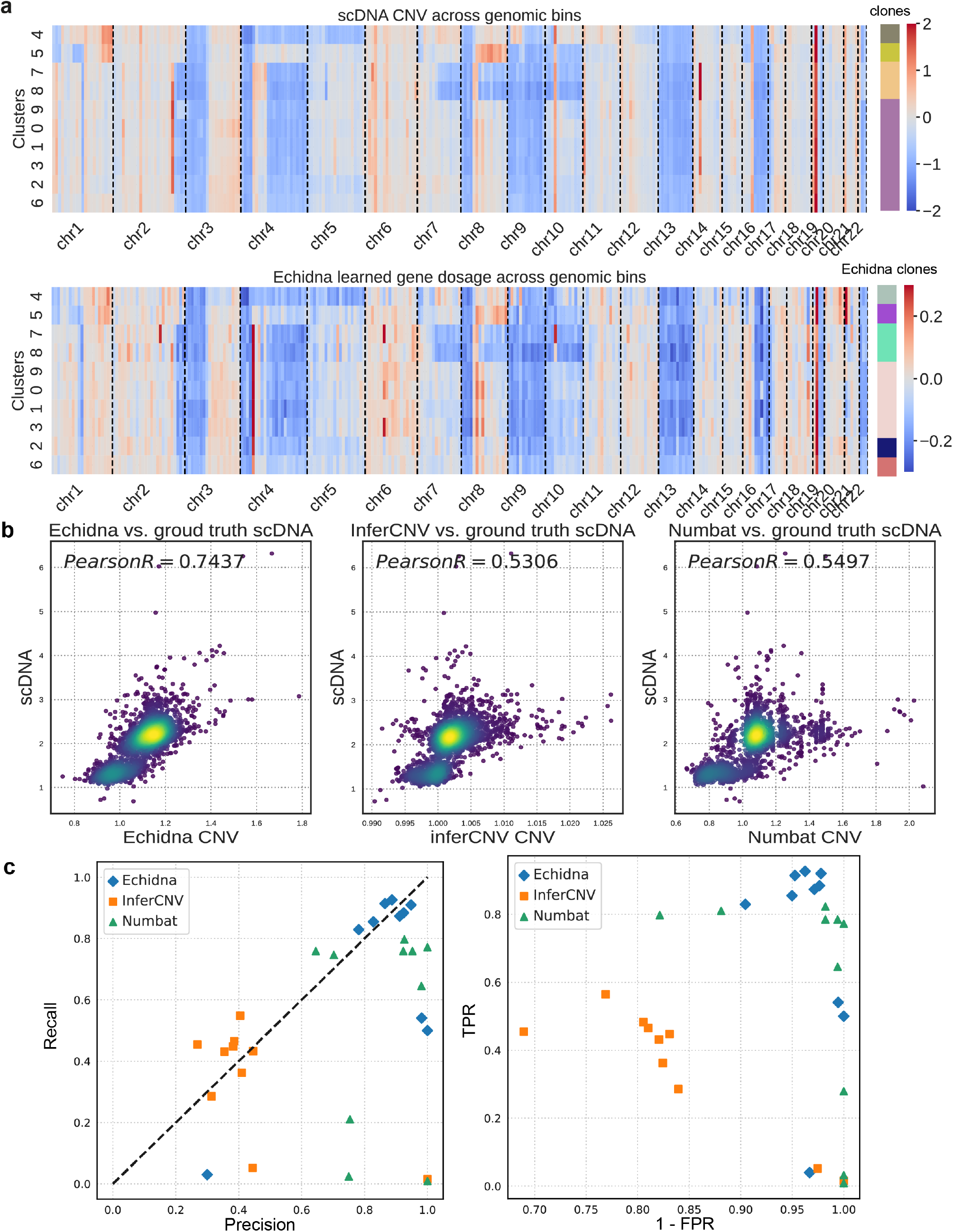
Echidna accurately estimates cluster-level CNA states and gene dosage. **(a)** Cluster-level CNA profile from an endometrial cancer sample (tumor 2 from [49]) showing ground truth scDNA collected via hybrid snDNA-seq (top) and inferred from hybrid scRNA-seq and matched WGS by Echidna (bottom). **(b)** Scatter plot comparing estimated CNA profiles by Echidna, InferCNV, and Numbat to scDNA data. Each dot represents a bin in a cluster and is colored by density. **(c)** CNA state prediction performance by different methods. Each dot represents a cluster (phenotypic state).

**Figure S3:**
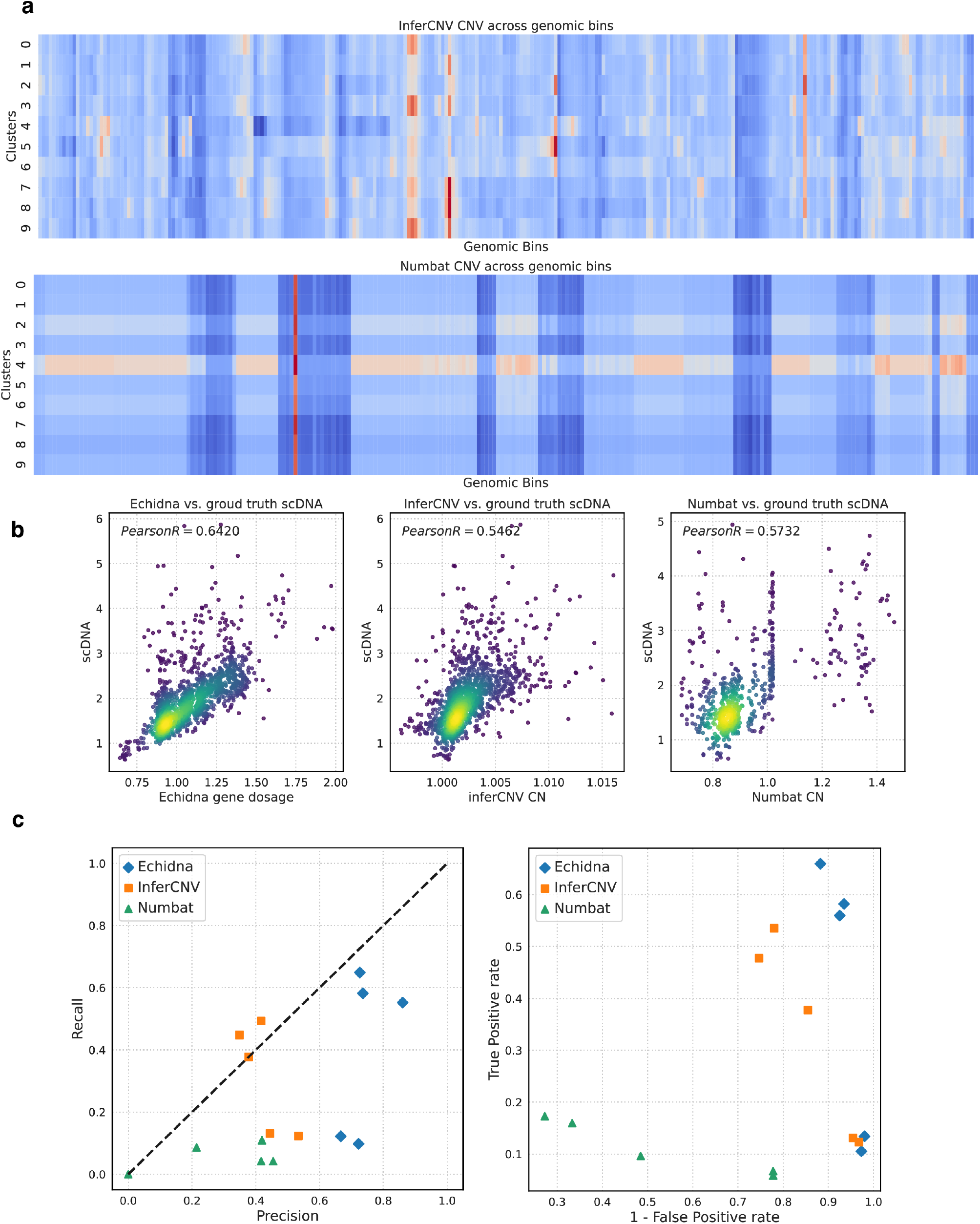
Benchmarking CNA profile estimation and CNV state prediction on an independent endometrial cancer sample. **(a):** Cluster-level CNA profile inferred by InferCNV and Numbat for tumor 5 from [49]). (b): Scatter plot comparing estimated CNA profiles by Echidna, InferCNV, and Numbat to scDNA data. Each dot represents a bin in a cluster and is colored by density. (c): CNV state prediction performance by different methods. Each dot represents a cluster (phenotypic state) in the tumor sample.

To evaluate the CNV state prediction performance within each cell cluster, we determined ground truth state labels by applying Gaussian HMMs to the scDNA CNA profiles across genomic bins. Each bin was categorized as amplification, deletion, or neutral based on the emitted HMM states. A separate HMM model was fitted to the CNA profile of each cell cluster. We obtain Echidna’s predicted CNV states by the same post processing steps mentioned in the overview. We assign states to the cluster-level CNA profiles estimated by InferCNV by applying thresholds based on the standard deviations within each cluster. Numbat CNV states were acquired directly from the MAP estimates. We calculated precision, recall, true positive rate, and false positive rate to systematically assess CNV state prediction performance for each cell cluster. We observe superior performance by Echidna in predicting CNV states (mean F1 score = 0.7456 across clusters) in most clusters compared to InferCNV (mean F1 score = 0.3541 across clusters) and Numbat (mean F1 score = 0.6604 across clusters; **Fig. 3c, left**; see **Methods**). Echidna also achieves higher true positive rates while maintaining low false positive rates compared to InferCNV and Numbat (**Fig. 3c, right**).

Altogether, these results highlight the advantage of integrating scRNA with WGS data, where the former ensures accountability for phenotypic variability, and the latter serves as a constraint on cluster-specific CNA inference. It also demonstrates the strength of linking gene dosage to expression within a unified model.

### 2.4 Echidna tracks clonal evolution using longitudinal data

Phenotypic plasticity plays a major role in the evolution of normal and pathological systems. To understand the extent to which transcriptional variability over time is enabled by gene dosage, Echidna integrates scRNA-seq and WGS data from different time points in the same system, thus linking clonal evolution to the temporal dynamics of gene expression. Establishing a strong link between genotype and phenotypic plasticity can characterize the cancer cell-intrinsic features that are critical for treatment response or disease progression. For example, applying Echidna to longitudinal data collected before and after anti-cancer treatment can explain if tumor cells respond or resist therapy because of clone-specific CNA events, or other factors such as epigenetic variability or involvement from extrinsic sources from the microenvironment such as modulated immune activity.

To demonstrate Echidna’s utility and impact for studying temporal evolution, we applied Echidna to our previously published data from a melanoma patient treated with anti-PD-1 immune checkpoint blockade [20]. Single-cell transcriptomic data from tumor cells exhibit pronounced heterogeneity between the pre-therapy and on-therapy specimens (with little overlap in clusters; **Fig. 4a**). This heterogeneity is not due to technical (batch) effects, as shared immune cell states were detected between the time points[20]. We first confirmed that Echidna accurately reconstructs both scRNA-seq and WGS data across the two time points (**Fig. 4b,c**), showing that the multitimepoint model effectively fits the data.

**Figure 4:**
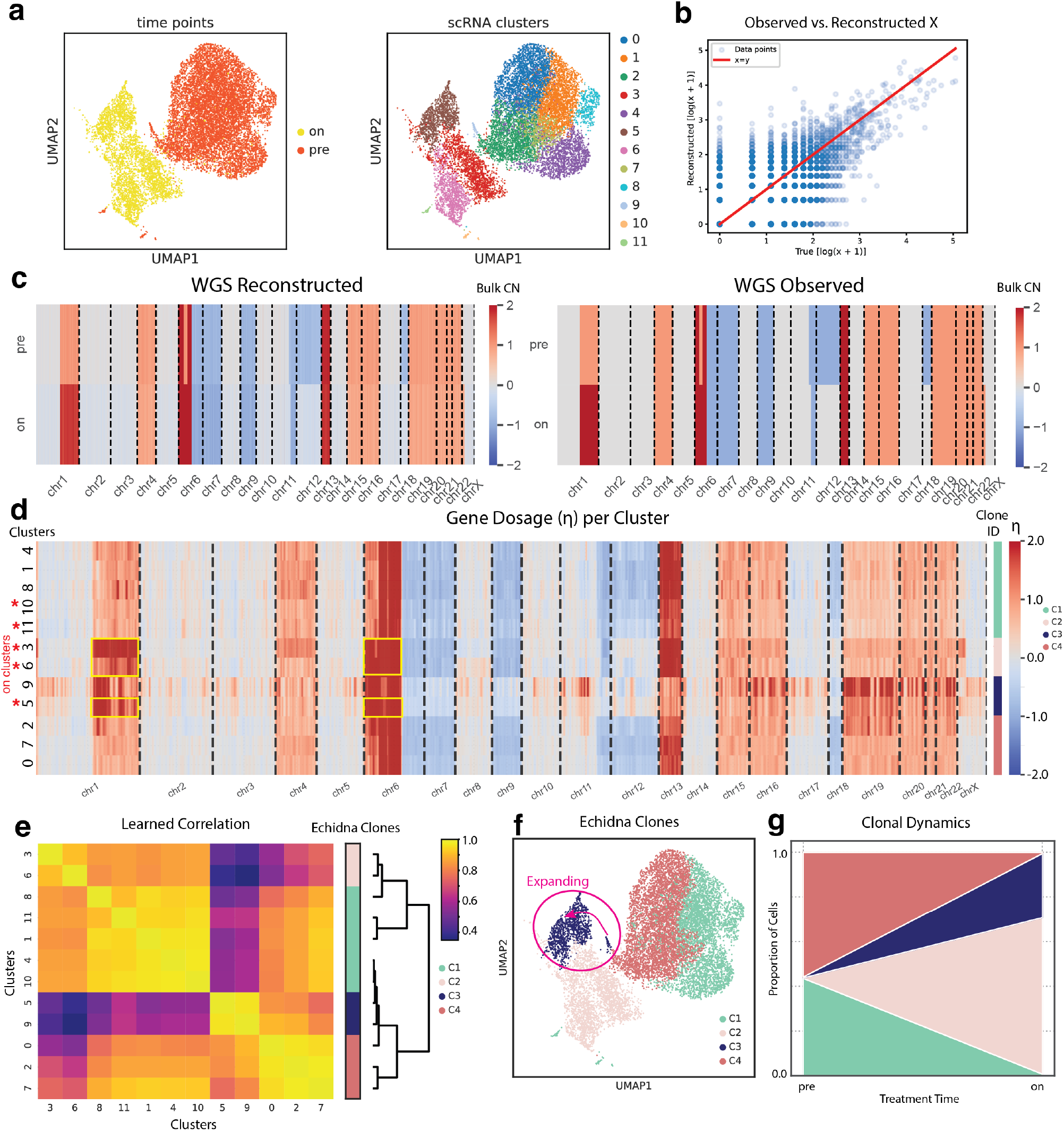
Echidna applied to a multi-time point dataset from a melanoma patient. **(a)** UMAP projection of scRNA-seq of tumor cells combined across samples, colored by treatment timepoint: pre- and on-immunotherapy (left) and phenotypic (Leiden) clusters (right). **(b)** Observed vs. model reconstructed scRNA-seq data. **(c)** Reconstructed (left) and observed (right) WGS data for each treatment time point. **(d)** Echidna inferred gene dosage values per cluster. Red stars denote clusters in the on-treatment time point, while the right bar groups clusters into clones. Copy number states are shown in **Fig. S1b. (e)** Inferred correlation matrix from Echidna, with clusters grouped by clonal structure. **(f)** UMAP projection of scRNA-seq of tumor cells combined across both data points, colored by the clonal structure revealed through correlation. **g** Proportion of cells belonging to each Echidna clone at each time point.

**Figure S4:**
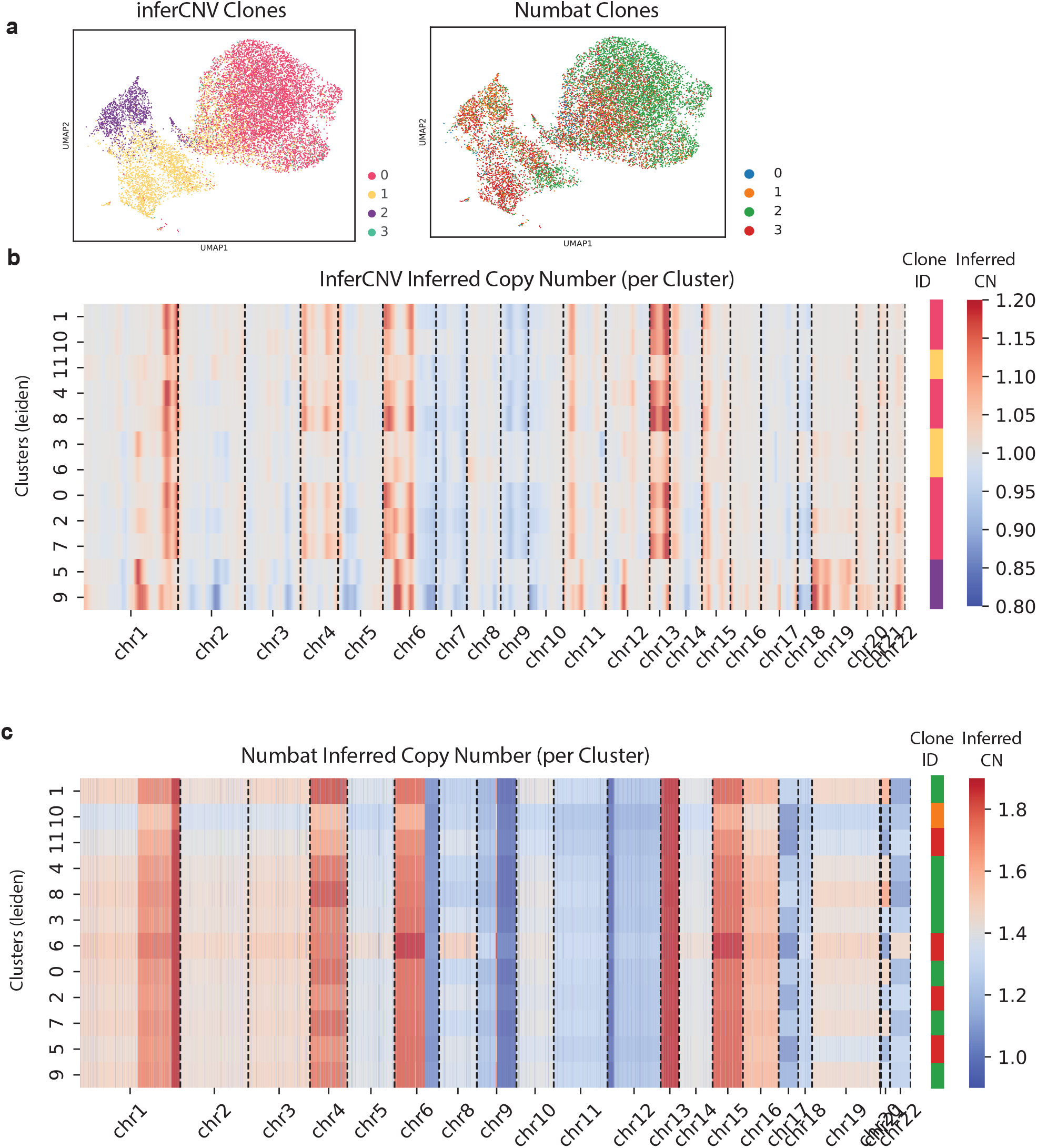
Performance of other methods on melanoma data. **(a)** UMAP projection of scRNA-seq of tumor cells combined across two treatment timepoints similar to **Fig. 4a** with cells colored by clones defined from inferCNV results (left) and Numbat clones (right). **(b)** InferCNV-inferred copy number averaged across cells for each phenotypic cluster. **(c)** Numbat-inferred MAP ratio to diploid averaged across cells for each phenotypic cluster.

**Figure S5:**
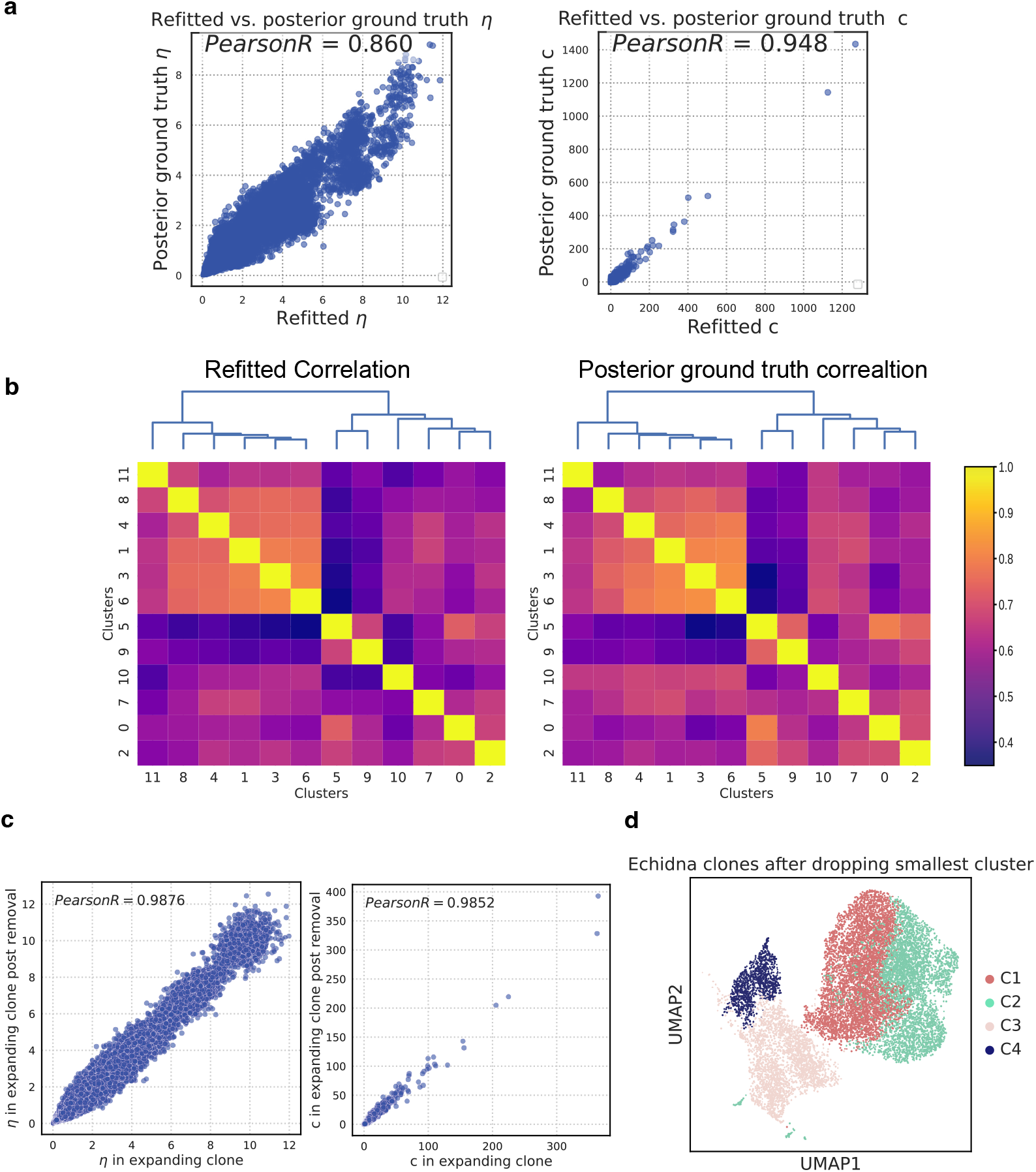
Performance on simulated data and robustness analysis in melanoma data. **(a)**: Comparing inferred gene dosage (*η*) and transcription rate(*c*) on semi-synthetic data to posterior ground truth. **(b)**: Comparing inferred correlation matrix and its structure on semi-synthetic data (left) to ground truth (right). Correlation matrices are obtained by normalizing the learned/ground truth covariance matrix parameters. **(c)**: Gene dosage and transcription rate in the expanding clone before and after removing the noisy cluster within the clone. **(d)**: Inferred clones from the melanoma two time point dataset (**Fig. 4**) after removing the smallest cluster showing robust inference of clonal structure.

We then examined inferred cluster-specific CNAs, showing diverse patterns of amplification and deletion (**Fig. 4d; Fig. S1b**). We find that clusters with similar CNAs are grouped together, defining clonal identities (**Fig. 4e**). Importantly, Echidna uncovers the clone shared between the two time points (clone 3), that was previously reported in this patient (**Fig. 4f**)[20]. This clone, marked by amplifications in chromosomes 1 and 6, is detected in a small subpopulation of cells pre-treatment (cluster 9) and grows in proportion post-treatment (cluster 5; **Fig. 4g**). Clone 2, which shares these amplifications, is additionally only present in the on-treatment time point, suggesting potentially shared resistance mechanisms. In contrast, clones 1 and 4 exhibit responsive dynamics, i.e. contracting with therapy. We compared the inferred CNAs and clonal structures from Echidna to other methods and found that while inferCNV was able to correctly identify the expanding clone, Numbat did not find the meaningful differences between clusters and therefore did not successfully recover the clonal structure (**Fig. S4**).

To further assess Echidna’s performance and robustness in recovering clonal architecture from time-series data, we applied it to semi-synthetic data. Specifically, we used inferred model parameters from fitting to the melanoma data and simulated new data from Echidna’s generative process. We thus have the ground truth parameters for this semi-synthetic data that has a realistic distribution. We show accurate reconstruction of gene dosage(*η*), transcriptional rate(*c*), and clonal architecture and evolution(covariance matrix) (**Fig. S5a,b**).In reality, very small or noisy clusters frequently appears in scRNA-seq data. These outlier cells often have noisy transcriptional profiles and can disturb correlation structures between clusters due to low sample size and high variance.To display the model’s robustness to these small or noisy clusters, we show robust recovery of both gene dosage and transcriptional rate in the expanding clone by removing the cluster with the lowest frequency in the melanoma patient in one time point and re-fitting the model (**Fig. S5c**). Additionally, we show the robustness of inferred clonal evolution structure (**Fig. S5d**). Overall, these results show Echidna provides a powerful framework for analyzing longitudinal data, uncovering clonal evolution, and enabling new insights into its impact on the temporal dynamics of phenotypic states in both real and simulated systems.

### 2.5 Echidna quantifies the effect of gene dosage on temporal gene expression dynamics and phenotypic plasticity

Our findings in the melanoma case study showcase an example of differential responses between clones within a tumor sample, further motivating approaches for disentangling the effect of genetic features on clone evolution. Since Echidna’s generative process explicitly encodes dependency between the transcriptional rate unique to a cluster (phenotypic state) and its CNAs, we can directly leverage the inferred posterior parameters to quantify this effect. Specifically, we compute the proportion of variance in the transcriptional rate of each gene *i* in cluster *k* (according to the inferred posterior distribution of *c*_*ik*_) that is explained by the dosage of that gene (*η*_*i*_) in its corresponding clone (see **Methods**). This metric thus provides a quantitative measure of *gene dosage effect (GDX)* for every gene in each cluster. Overall, we observe higher GDX values in genes located in amplified or deleted regions as expected (**Fig. S1d**). However, the relationship between gene dosage and GDX is variable across heterogeneous phenotypic states (**Fig. S1d**).

We utilize this metric to gain deeper insight into mechanisms leading to clone resistance in the melanoma tumor sample. While most approaches aim to identify features predictive of treatment response at the sample (e.g. patient) level [5, 24, 39, 50–52], Echidna allows for the study of the drivers of phenotypic plasticity at the clone level, truly leveraging the heterogeneity profiled with scRNA-seq and linking it to genetic features. We dissect the melanoma tumor into clones that expand (i.e. increase in proportion) with treatment and those that contract (decrease in proportion; **Fig. 4g**). We then focus our investigation on genes that are differentially expressed (DE) between the expanding and contracting clones to identify drivers of heterogeneous response. Visualization of the genomic positions and inferred gene dosage reveals that a subset of DE genes are concentrated in regions of genomic amplification (**Fig. 5a, b**). Thus, by scanning genomic regions predicted to have an increase in copy number while encompassing a high density of DE genes (**Fig. S1b, c**), we identify *“hotspots”* of gene activity corresponding to phenotypic differences between clones (**Fig. 5a,b**). The GDX metric then allows us to verify if these hotspots are driven by underlying genomic alterations. This approach places the DE genes unique to clones predictive of response or resistance to treatment in the context of genomic alterations and quantifies their association through GDX scores.

**Figure 5:**
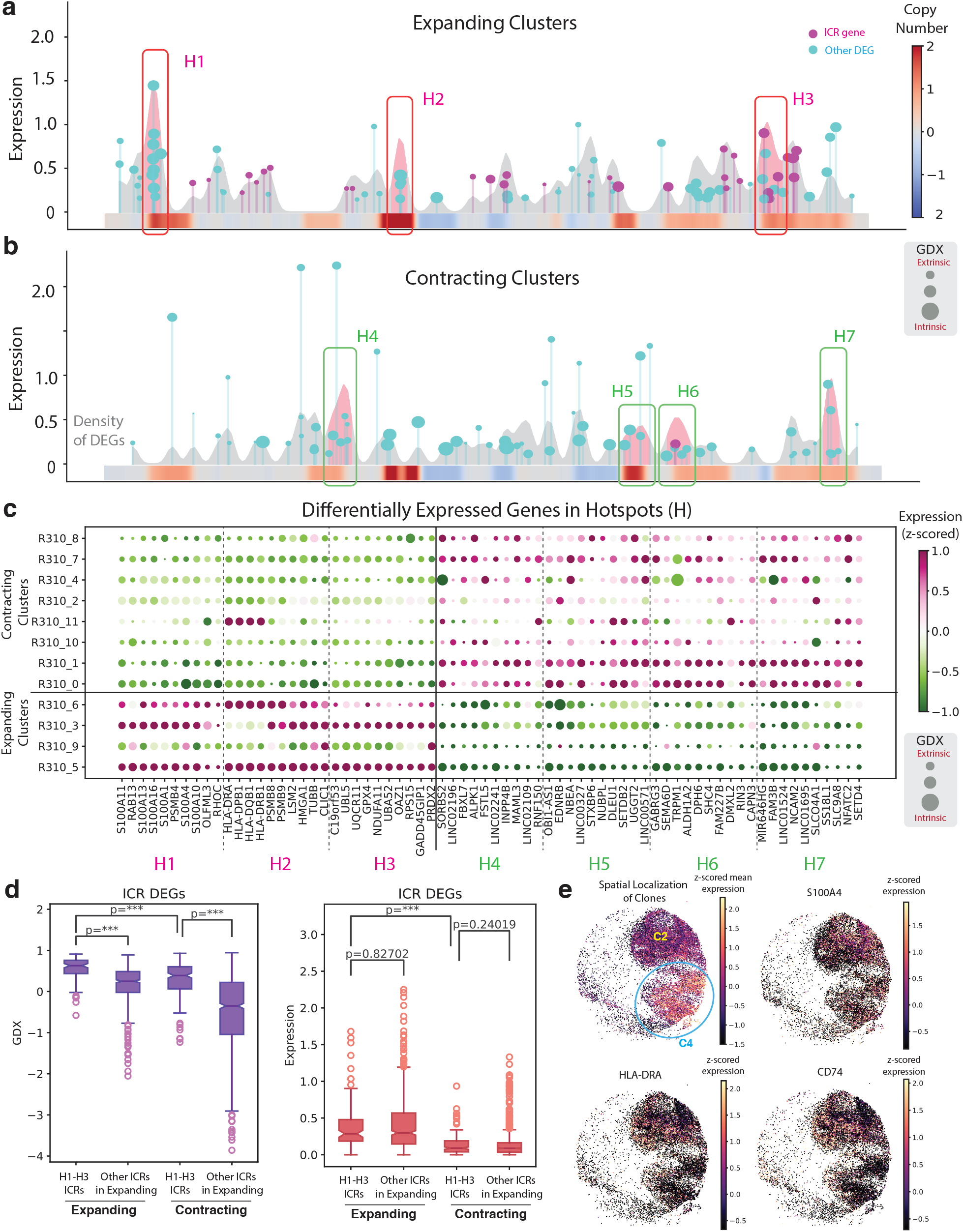
Gene Dosage Effect (GDX) reveals mechanisms underlying clone-specific response to treatment. **(a-b)** Top differentially expressed (DE) genes in clusters expanding **(a)** and contracting **(b)** with treatment, visualized as lollipops with x-axis corresponding to genomic position. Density plot reflects the distribution of DE genes by genomic position. Height of each lollipop corresponds to the expression of each gene, while dot size reflects the absolute value of the GDX, such that larger dots denote stronger dependency of expression on gene dosage. Mean gene dosage is plotted underneath as a heatmap. ICR genes are highlighted in pink. **(c)** Dotplot of top DE genes in the hotspots highlighted in (a,b) for each cluster. Dot size corresponds to the absolute value of GDX, while color represents z-scored expression. **(d)** GDX values (left) and expression (right) for ICR genes upregulated in expanding clones (resistant to treatment) that are located in genomic hotspots H1-H3, and those not in any hotspots. P-values were computed using the Mann-Whitney U-test. ^***^ denotes *p <* 1*e* − 4. **(e)** Spatial transcriptomic (Slideseq) data for the sample collected at the on-treatment timepoint. Spots are colored by the expression of clone 4 signature (also circled) with the location of clone 2 labeled, and expression of key genes with high spatial variation.

Applying this analysis across expanding and contracting tumor clones within the melanoma sample enables the characterization of clone-specific response mechanisms. We first confirmed that the top DE genes in the detected genomic hotspots had significantly higher GDX (*p <* 1*e* − 3 for all clusters) compared to non-hotspot DE genes, verifying that the increased expression in hotspots was due to underlying amplifications. In the contracting clones, we identified four separate hotspot regions **(Fig. 5b, c)**. Top DE genes of interest in these regions encode for two families of molecules, zinc finger proteins (e.g., *ZFP64, ZNF208, ZNF91, ZNF790-AS1, ZNF506, ZNF875, ZNF529, ZNF146*) and long intergenic non-coding RNAs (e.g., *LINC02438, LINC02267, LINC01531, LINC00906, LINC01524*). The genes we uncovered from these families are understudied in melanoma, but other genes from these families have known mechanistic involvement in development, progression, and treatment response in melanoma [53, 54] and other cancers [55], and have been found to be relevant predictors of immunotherapy response in melanoma [56, 57]. For example, lower levels of RNA editing in nucleotide-binding genes *ZNF124, 490*, and *827* predict immunotherapy sensitivity in pembrolizumab-treated patients, and the same for *ZNF226, 329, 426* and *836* in nivolumab-treated patients [56]. High expression of TRIM28, a cofactor for zinc finger proteins, corresponds to higher melanoma stemness and depleted immune infiltration [54]. Several long non-coding RNAs suppress many melanoma processes, including metastasis and proliferation (e.g., “sponging” of tumor suppressor gene miRNAs by *LINC00459* or *LINC00961*), phenotype switching (e.g., transcriptional regulation via *TINCR* expression), immune evasion (e.g., *LIMIT* expression transcriptionally inducing tumor immunogenicity), and drug resistance (e.g., *TSLNC8* expression abrogating MAPK signaling via PP1*α* binding) [53, 57]. In summary, GDX scores are a powerful metric to investigate mechanisms of clone-specific response to treatment.

Among the hotspots in the expanding clones, we identify H1, (**Fig. 5a, c**) which contains multiple S100A family genes. The role of this family of genes in melanoma as early diagnostic markers as well as putative drivers of tumor invasion has been well established [58, 59], and falls in line with their cell-intrinsic effect predicted by GDX. We also identified a second hotspot, H2, s that was highly enriched in MHC-II genes. This finding reflects recent discussions on the complex role of MHC-II in melanoma response to immunotherapy, where MHC-II genes were hypothesized to function as a marker of tumor inflammation, and alone were not sufficient for the induction of an immune response [60]. Our results demonstrate that these genes are not only found to be upregulated in resistant clones, but they are also significantly linked with underlying genomic amplifications.

We further note the increased presence of genes from the immune checkpoint resistance (ICR) gene signature [20] in expanding compared to contracting clones. A subset of ICR genes were found to be present in hotspot regions (H3), and others scattered across the genome. This combination suggests that in addition to genomic changes, there are additional extrinsic factors at play in tumor resistance to immunotherapy, which may include aberrant tumor-immune cell interactions and local environmental effects. We demonstrate that our GDX score is able to disentangle intrinsically driven ICR genes from those unlinked to genomic events. Specifically, we define two sets of genes for statistical testing: ICR genes upregulated in expanding clones that are localized to hotspot regions (H1-H3), and non-hotspot ICR genes upregulated in expanding clones. These two gene sets were then examined in both expanding and contracting clusters. We show that the GDX is significantly different not only between expanding and contracting clusters, but also between hotspot and non-hotspot genes within categories. Conversely, gene expression only captures the difference between expanding and contracting clones (**Fig. 5d**). The increased granularity that can be discerned with the GDX metric may also be seen at the level of gene signatures, where enrichment scores computed for each signature reveal differences in invasion, antigen presentation, and immune resistance signatures; whereas expression only shows a difference in antigen presentation **(Fig. S6)**.

Finally, we interpret our GDX metric in the spatial context using previously published spatial transcriptomic (SlideSeq) data [20] for the on-treatment timepoint. We first highlight that clones identified by Echidna appear as spatially distinct regions in the tumor (**Fig. 5e**). Specifically, clone 4 which expresses the ICR signature [20] is spatially separated from clone 2, suggesting that clonal expansion occurred in a spatially segregated fashion following genetic alterations in early founder cells. We then computed the Moran’s I spatial autocorrelation score, which was adapted for use in spatial transcriptomics by [61], to find genes with distinct spatial localization among growing clones. Among the top genes ranked by the Moran’s I score, we find *S100A4* (GDX=0.59), a gene known to promote tumor metastasis through epithelial mesenchymal transition (EMT) [62], and HLA-DRA (GDX=0.4), an MHC class II family gene – with the high GDX scores suggesting that genetic events drive spatial variability. In contrast, high Moran’s I and low GDX suggest extrinsic factors driving spatially varying expression, e.g. in *CD74* (0.11), which has been found to promote melanoma progression through activation of *AKT* signaling and binding to *MIF* [63] (**Fig. 5e**). Our method thus allows for the linking of distinct spatial patterns to underlying genomic alteration events. These results further suggest that Echidna may unveil underlying dynamics governing the transformation and localization of complex cell lineages.

## 3 Discussion

Phenotypic plasticity has emerged as an important concept in tumorigenesis, tumor progression, and resistance to targeted and immune-based treatments in multiple cancer types as well as other disease and normal contexts. While technical development readily enables the measurement ofsingle-cell gene expression and has uncovered transcriptional heterogeneity which may represent plasticity, a major challenge in the field has been a lack of methods that account for gene dosage. Understanding whether transcriptional changes are merely a consequence of gene dosage differences between cancer clones within a sample or across patient cohorts, and separating this effect from putative plasticity induced through interactions with other cells of the TME or non-genomically encoded events, remains a major unmet need in the analyses of tumor data.

While concurrent measurement of single-cell DNA and RNA-sequencing could address some of these challenges, coupling these methods remains a technically challenging task and is difficult to scale. In contrast, obtaining bulk WGS and scRNA-seq from the same specimen is feasible and scalable now. We therefore reasoned that a pragmatic solution to this challenge is integrative modeling of these two data modalities through devising a novel computational framework – Echidna - that uncouples gene dosage effect from extrinsic impact on plasticity, at scale, and over time, within and across patient samples.

The conceptual core of Echidna lies in its hierarchical modeling of the relationship between clonal identity and transcriptional programs, enabling a fine-grained dissection of cellular adaptation and clonal dynamics when data from multiple time points are available. Importantly, Echidna addresses a key gap in the field by quantifying the gene dosage effect (GDX), offering a direct measure of how CNAs influence transcriptional states within specific clones. In the context of cancer research, where clonal diversity often drives therapy resistance and relapse, Echidna provides a crucial tool for studying these processes at high resolution, and in a technically and computationally scalable format. In our application to melanoma, we identified a treatment-resistant clone that expanded following anti-PD1 therapy, marked by distinct CNAs and transcriptional reprogramming. Given the emerging recognition that large chromosomal aberrations rather than specific point mutations may determine drug resistance in different disease and treatment contexts[64], Echidna provides a framework to begin unraveling underlying mechanisms while accounting for different sources of plasticity. Beyond its ability to uncouple GDX from extrinsic determinants of plasticity, Echidna outperforms existing and widely used methods for the inference of CNAs in single-cell RNA-seq data.

In its current form, Echidna is applicable to cancer types where CNAs are a defining hallmark. However, this model can be extended for application to other genetic alterations including point mutations or epigenetic changes. For instance, SNVs can be detected from WGS data and built into the prior for cluster similarities. Similarly, other data modalities such as ATAC-seq can inform these similarities. The flexible multi-view generative process can also be expanded to integrate more than two data modalities in future work.

Beyond its immediate application to cancer, Echidna has broader implications for studying adaptation and evolution across biological systems. For instance, it can be adapted to study clonal dynamics in the immune system, where genetic variability and transcriptional plasticity influence responses to infections or vaccinations. Similarly, its application in developmental and regenerative biology could provide insights into how genetic alterations influence tissue repair or stem cell differentiation. Overall, Echidna represents a significant step forward in modeling the dynamic interplay between genetic features and phenotypic plasticity, with broad implications in cancer and beyond.

**Figure S6:**
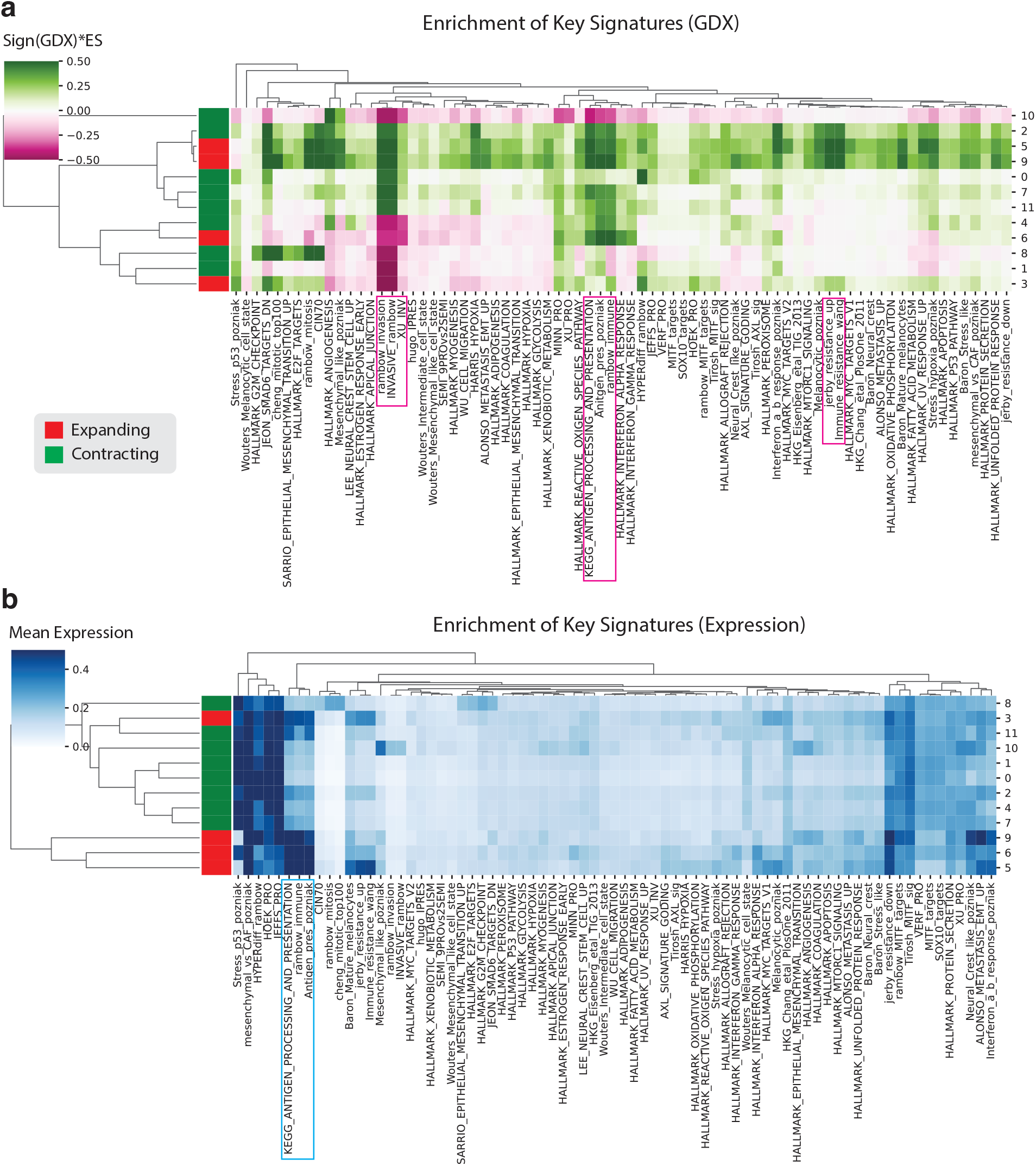
Expression and GDX-based enrichment of key gene signatures. **(a)** Enrichment of key signatures according to a permutation test applied on GDX scores for genes in a signature compared to other genes. Color corresponds to the sign of the GDX (indicating intrinsic vs extrinsic effect) multiplied by the enrichment score. **(b)** Enrichment of key signatures by gene expression

## 4 Acknowledgements

We thank Siran Li and Michael Wigler for helpful data sharing and discussion. We also thank Andrea Califano, Peter Sims, Yinuo Jin, and Aymeric Degroote for helpful discussions and feedback. J.L.F. acknowledges support from the Columbia University Van C. Mow fellowship and the Columbia Avanessians fellowship. K.H.H. was supported by the National Science Foundation Graduate Research Fellowship under Grant No. DGE-2036197. B.I. is supported by National Institute of Health (NIH) grants, R37CA258829, R01CA280414, the Pershing Square Sohn Cancer Research Alliance Award, the Burroughs Wellcome Fund Career Award for Medical Scientists; a Tara Miller Melanoma Research Alliance Young Investigator Award; the Louis V. Gerstner, Jr. Scholars Program; and the V Foundation Scholars Award. B.I. is a CRI Lloyd J. Old STAR (CRI5579). B.I. is supported by the Herbert Irving Comprehensive Cancer Center Human Tissue Immunology and Immunotherapy Initiative. B.I. and E.A. acknowledge support from NIH NCI grant R01CA266446, Columbia Research Initiatives in Science & Engineering (RISE) Award, and a pilot award from Herbert Irving Comprehensive Cancer Center (HICCC) NIH NCI grant U54CA209997. E.A. is supported by the NIH NCI grant R00CA230195, NIH NHGRI grant R01HG012875, R21HG012639, and grant number 2022-253560 from the Chan Zuckerberg Initiative DAF, an advised fund of Silicon Valley Community Foundation.

## 5 Author contributions

E.A. and B.I. conceived the study and provided overall supervision of the study. J.L.F, M.Z., W.O.B, J.D.M., N.B.V, A.P., I.A., and E.A. designed and developed the Echidna framework. J.L.F, M.Z., W.O.B., J.D.M, and E.A. contributed to implementation. J.L.F., M.Z., W.O.B, J.D.M., S.C., and E.A. analyzed data. J.L.F., M.Z., J.D.M., J.C.M., E.D.S., S.T., K.H.H., B.I and E.A. interpreted data. J.L.F, M.Z., K.H.H., B.I., and E.A. wrote the paper. All authors reviewed, contributed to and approved the paper.

## 7 Methods

### 7.1 Model specification

**Notations** Echidna is a Bayesian hierarchical model. We utilize variational inference to infer posterior distributions of latent random variables that generate observed variables through a designed generative process. Assume we have *T* time points, consider the WGS copy number vector 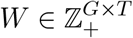 across *G* genes and the scRNA count matrix 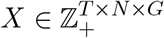 for *N* cells, where Z_+_ denotes a set of nonnegative integers. We denote the deconvolved gene dosage (copy number) in the *K* cell clusters as *η ∈*R^*G×K*^and the time-specific latent cell transcriptional rate as *c ∈*R_+_^*T ×G×K*^, where R is the set of real numbers and R_+_ is the set of positive real numbers. We also denote the proportion of observed cell clusters as *π ∈* R^*T ×K*^ and the cluster membership matrix as *z ∈* [0, 1]^*T ×N ×K*^. The libaray size (sum of counts per cell in scRNA-seq) is denoted as *l ∈* Z_+_^*T ×N*^. We denote *K* dimensional multivariate normal distributions as 𝒩 ^*K*^.

#### Generative process of Echidna

We build correlation structures between clusters of cells (within or across time points) informed by both scRNA and WGS. Clusters with highly correlated *η*s are interpreted as clones (which can be static and over time). Therefore, we treat the covariance matrix of *η* as a random variable. The lower Cholesky factors *L* of the covariance matrix Σ_*η*_ is given by:

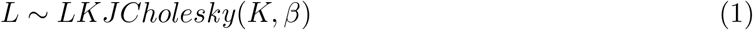

where *β* is the concentration parameter. For each cluster *k*, we sample cluster variances *σ*_*η*_ from inverse Gamma distributions as:

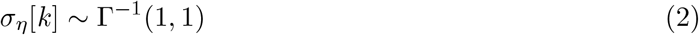

We combine the Cholesky factors and the diagonal variances to parametrize the full covariance of *η* as:

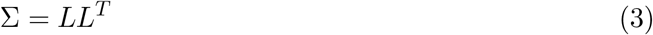

and updated as:

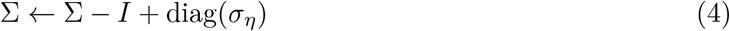

where we obtain the full covariance with unit variance in (3) and scale the diagonal elements with the sampled variance *σ*_*η*_ in (4). We sample the mean of *η* from a *K* dimensional multivariate normal centered around neutral copy number (2) such that:

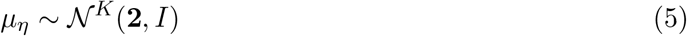

We model gene dosage as deconvolved copy number profiles across *K* clusters for all genes as *G* different *K* dimensional sampling distributions. The gene dosage of gene *i* is given by:

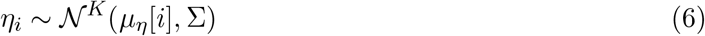

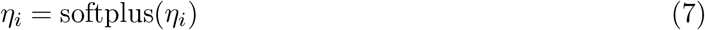

where in (7) we apply a softplus activation to the sampled gene dosage to keep it positive. Assuming gene dosage in a specific clone does not change dramatically over time, the observed copy number *W*_*i*_ for gene *i* at time point *t* is in turn parameterized as a mixture of *K* clusters weighted by cell proportions at the time point *π*_*t*_ as:

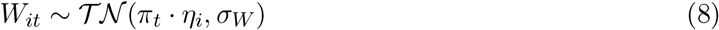

Contrary to the gene dosage, we expect transcription profiles of cells to change across time points leading to phenotypic plasticity. By default we set *σ*_*W*_ = 0.05 to account for the gene-specific variability in copy numbers. For gene *i*, cluster *k*, and time point *t*, we model the transcription rate of cells in cluster *k* as:

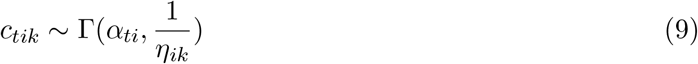

where the gene dosage *η* is the scale parameter of the Gamma distribution. The shape is given by a free parameter *α*_*ti*_. As a result, we establish a hierarchical structure in which the gene dosage impacts both mean and variance of the transcriptional rate. Finally, to obtain the single-cell RNA counts *X*, we identify the cluster that each cell belongs to and the corresponding transcription rate vector. We also account for the effect of library size and cluster-specific bias in gene dosage. We sample *X* at time point *t* as:

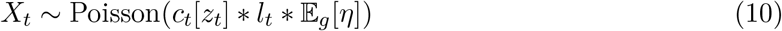

where *z*_*t*_ is the cluster membership matrix at time point *t*.

### 7.2 Variational inference

#### Variational family for lower cholesky factors

We first construct the variational distribution of the lower Cholesky factors(*L*) of the covariance of gene dosage(*η*). Formally, we recover the lower Cholesky factor of a *K × K* matrix, denoted as *L*. We need 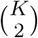 free parameters which we denote as *y*. We assume the free parameters follow a diagonal normal distribution with prior 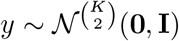. We learn the means and the variances of *y* as free parameters. From *y*, we can construct latent lower triangular matrix *ζ ∈* R^*K×K*^ by filling the below diagonal entries by row with transformed *y*:

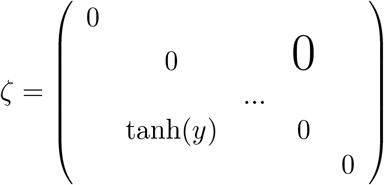

Then we can recover the cholesky factor *L* as:

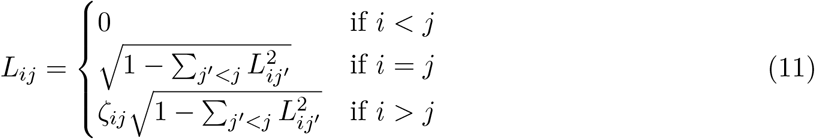

Denote the function mapping from *y* to *L* as 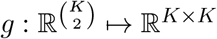, the variational distribution of *L* is given by:

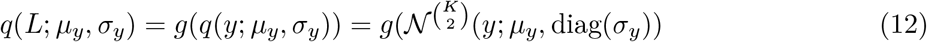

#### General variational family structure

For notation simplicity we drop dependency on observed random variables. The variational family of Echidna is given by:

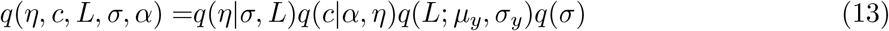

which can be factorized into:

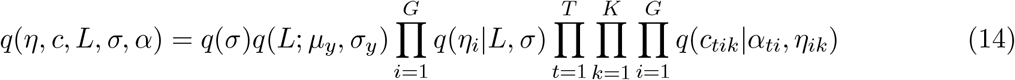

Mean-field variational inference is applied to learn posterior for each latent random variable. We parameterize the variational distributions as:

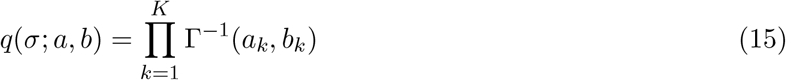

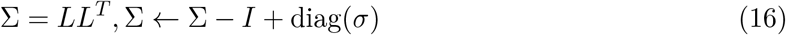

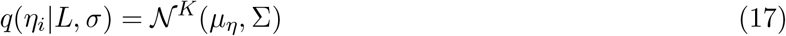

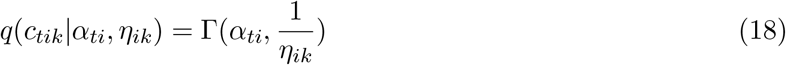

where 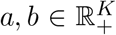 are variational parameters for the inverse gamma distribution on *σ*. Notice that although the notations of variational parameters remain unchanged for clarity, these are different parameters compared to the parameters in the generative process.

#### The variational inference objective

Variational inference aims to minimize the KL divergence between the variational posterior and the true posterior, which is equivalent to maximizing evidence lower bound. We will denote the variational family as *q*_*θ*_ where *θ* is the set of all variational parameters and generative densities as *p*_*ϕ*_ where *ϕ* is the set of all generative parameters. The evidence lower bound is given by:

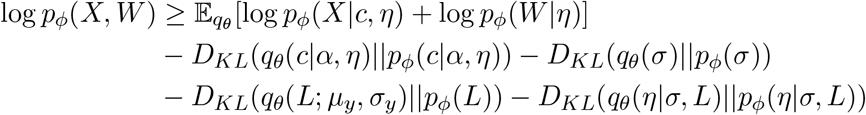

Again, we note that parameters of *q*_*θ*_ and *p*_*ϕ*_ are different sets of parameters despite the identical notations.

### 7.3 GMM model to determine cluster-specific neutral CNA

To estimate neutral CNA within each cluster from the learned gene dosage(*η*), we fit a 1-d Gaussian Mixture Model (GMM) to account for the different levels of CNA signal. More specifically, for the *k*-th cluster, the gene dosage vector is given by 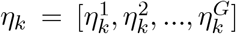. With *P* components, the probability density function of 1d GMMs are given by:

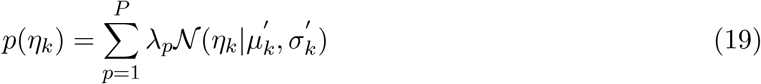

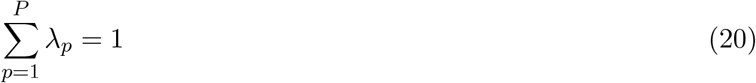

The model is fitted with an Expectation-Maximization(EM) algorithm and we use the output to predict which component each 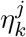 can be sampled from. We then estimate the neutral CNA for cluster *k*, denoted as 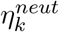 as:

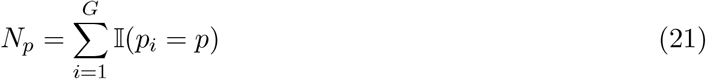

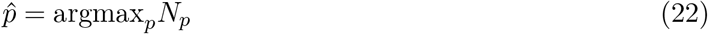

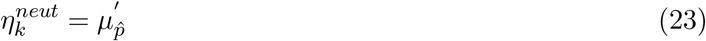

We do this for every cluster and construct vector 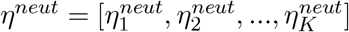. We can then shift the values of *η* by *η*^*neut*^ to center neutral CNV at 0 for all clusters.

### 7.4 HMM model for state prediction

To predict CNV states from the inferred gene dosage values (*η*), we first incorporate genomic positional information by smoothing the posterior estimate of *η* with a 1-d Gaussian filter per cluster along the genome. This approach accounts for gene level noise and enhances sequential dependency of neighboring genes. We then fit GMMs to the smoothed copy number estimates for each cluster, and adjust *η* so that neutral CNA is centered around zero (see **GMM model to determine cluster-specific neutral CNA**). We assume that the most frequent GMM component is the neutral state. We acknowledge that this assumption may not be valid in cases of multiploidy, in which case this model may require additional supervision. With the smoothed and adjusted *η*, we fit a HMM with Gaussian emission probability for each cluster-specific gene dosage vector. More specifically, consider smoothed and adjusted gene dosage *η*_*k*_ in cluster *k*, the joint likelihood of *η*_*k*_ and the sequence of hidden states *s* = [*s*_1_, *s*_2_, .., *s*_*G*_] is given by:

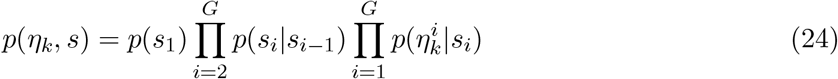

We fit the model with the Viterbi algorithm to find the state sequence *s*^*\**^ = argmax_*s*_*p*(*η*_*k*_, *s*) that maximizes the joint likelihood. This state sequence is the predicted CNV states along genes (or genomic bins).

### 7.5 Clone structure reconstruction

To reconstruct clone structure, we leverage the correlation structure across clusters to construct more robust hierarchical cluster trees. The correlation matrix can be obtained by normalizing the learned covariance parameter Σ. If the learned covariance matrix is noisy due to low sequencing depth or technical artifacts, we estimate the empirical correlation matrix from the smoothed learned gene dosage *η* to obtain a de-noised version. More specifically, small, spurious spikes or dips in correlation values are averaged out while pronounced correlation signals remain unchanged. To obtain the clone structure, we perform hierarchical clustering with Ward linkage with the covariance matrix of gene dosage. More specifically, the distance matrix for hierarchical clustering *D* is given by:

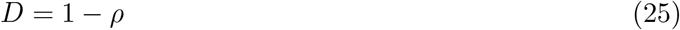

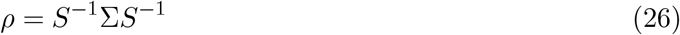

where *ρ* is the correlation matrix and *S* is the diagonal matrix of standard deviations(square root of the diagonal elements of Σ). If we choose to estimate empirical correlation, the correlation matrix *ρ* is given by:

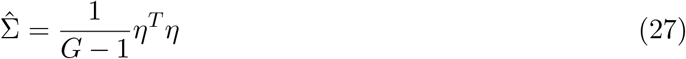

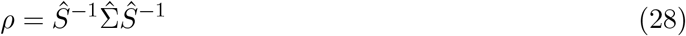

where *η* is the learned gene dosage,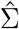 is the empirical covariance of *η* and *Ŝ* is the diagonal matrix of empirical standard deviations. We can then establish a threshold, either manually or based on the cophenetic distance of the hierarchical dendrogram, to derive flat clusters, which are subsequently identified as clones.

### 7.6 Gene Dosage Effect

To understand the relationship between gene dosage (*η*) and transcriptional rate (*c*), we quantify statistical dependency between the random variables in the model. Such a relationship can be biologically interpreted as gene dosage effect (GDX) which reflects how much gene expression changes depending on genetic level alterations. Genes whose expression changes can be well correlated with genetic variations such as copy number, i.e high dosage effect, can be interpreted as intrinsic programs. Conversely, genes whose changes in expression do not depend on genetic variations, i.e. low dosage effect are likely to be driven by extrinsic or non-genetic effects. To quantify such a relationship, we develop the following statistical framework. Consider the learned posterior distribution of *c* in one time point, we can factorize the densities to *K* different densities in each cluster *k*:

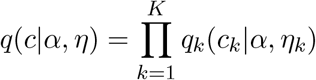

We then sample from the cluster-specific posterior distributions and get samples of *η* and *c* for all genes local to cell clusters. These samples live in R^*G*^ so they are per-gene measurements. We then compute the variance explained metric as the gene dosage effect (GDX) *s*_*k*_ in cluster *k* across these posterior samples to quantify dependencies between *η* and *c* for each gene for a specific phenotypic state *k*:

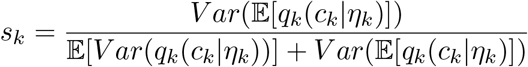

This metric allows us to score each gene in each subpopulation based on how its transcriptional variability is induced. We construct differential enrichment tests to capture the variation of plasticity-inducing mechanisms with gene signatures and connect these signatures to clinical outcomes such as clonal expansion or treatment response.

### 7.7 Gene signature enrichment based on gene dosage effect

To quantify gene signature enrichment and its statistical significance, we first rank all genes with the GDX score(for detail see **Gene Dosage Effect**). Denote the set of all ranked genes *G* and the gene signature *S ⊂ G*, we compute a enrichment score(ES) to reflect how *S* are distributed among the ranked set *G*. The enrichment score is the absolute maximum deviation of the running-sum scores, namely:

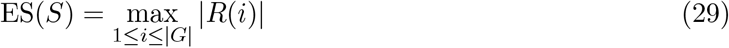

where |*G* | is the number of genes in the ranked gene set and *R*(*i*) is the running-sum score at position *i*, which is given by:

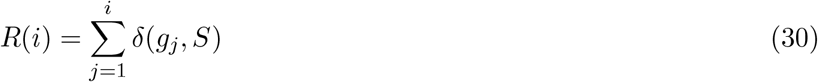

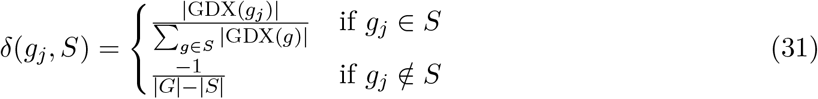

To access the statistical significance of enrichment score for gene signature *S*, we generate a null distribution by permuting the GDX scores and re-computing ES. The nominal p-value is then given by:

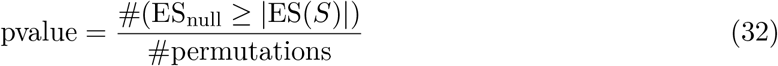

We lastly adjust the p values for multiple hypothesis testing with Benjamini-Hochberg correction.

### 7.8 Preprocessing of endometrial cancer data

Paired snDNA/snRNA data along with corresponding WGS were obtained from [49]. The WGS data were mapped from genomic regions to genes in order to fit the standard Echidna expected inputs. Similarly, for ground truth validation. the snDNA was mapped from the bins defined by [49] into individual genes. To run Echidna, only a few hyperparmeters have to be adjusted. We increased *σ*_*y*_ to 0.1 during variational inference to account for the relatively low cell count. We also changed the initialization of the mean of the prior of *µ*_*η*_ to be the mean of the mapped WGS data.

### 7.9 Benchmarking against other methods

#### Comparison to inferCNV and Numbat

InferCNV uses scRNA-seq data with reference cells that do not have CNV events to identify single-cell level copy number alterations [42, 65]. Numbat takes advantage of genotype information from DNA data and infers copy number alterations in a haplotype-aware fashion [43].

When running InferCNV on the tumor 2 dataset, we lowered the cutoff from 1 to 0.01 due to the sparseness of our count matrix and used default values for the rest of the parameters: cluster by groups, denoise, and HMM were all kept as true. For the Numbat run, the genome was changed to “hg19” and the number of cores was increased from 4 to 20 to speed up the run, but all other default parameters were left unchanged. We used T cells as our reference for both tools. For the tumor 5 dataset, we lowered the InferCNV cutoff even further to 0.001 because of the increased sparseness of the tumor 5 count matrix compared to that of tumor 2. We also increased the max entropy parameter in Numbat from 0.5 (default) to 0.75 to prevent all CNVs from being filtered out. All other parameters were kept the same as in the tumor 2 runs, and any non-tumor cells in the dataset were used as reference.

#### Benchmarking clone reconstruction

Echidna clones are called by performing Ward linkage based hierarchical clustering with the inferred covariance Σ_*η*_ (for details see **Clone structure reconstruction**). Numbat clones are directly obtained from the output of the last iteration of phylogeny optimization. InferCNV clones are identified by first pseudo-bulking the single-cell CNA estimates at the cluster level, followed by applying K-means clustering. We assign K as the ground truth number of clones identified from scDNA data. The performance metrics used are purity, adjusted mutual information, and adjusted rand index. Assume we have *N* cells and *K* clones. The purity is given by:

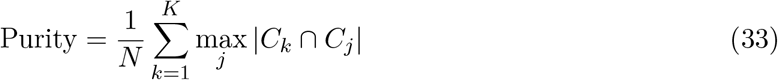

where *C*_*k*_ are cells belonging to clone *k* and *C*_*j*_ are cells belonging to ground truth clone *j*. This score is biased towards more clones, but measures the number of cells in the intersection of each clone *k* with ground truth clone *j*. To account for the number of clones bias, we introduce two metrics adjusted for chance. Given two sets of clone labels *U* and *V*, the adjusted mutual information(AMI) is given by:

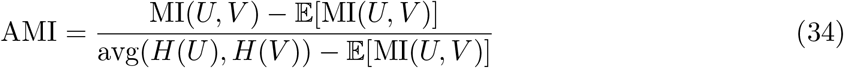

where *H*(.) is the entropy. The adjusted rand index(ARI) is given by:

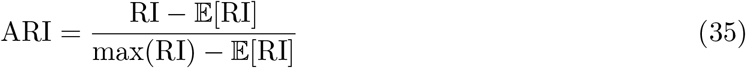

where RI is the ratio between the number of agreeing cell pairs and the total number of cell pairs. The pairs are constructed over identified clones and the ground truth clones.

#### Benchmarking CNA estimation

Since scDNA CNA profiles are divided into 10Mb bins, we apply the binning introduced in [49] to the estimated cluster-level CNA profiles from all methods. For Echidna, we directly used the learned gene dosage parameter *η*. For inferCNV, we pseudo-bulk the single-cell CNA estimates at the cluster level, with each cluster’s CNA profile defined as the mean of the CNA profiles of all cells within the cluster. For Numbat, we expanded the clone-level pseudo-bulked total copy number ratio relative to diploid, where the estimated CNA profile for each cluster corresponds to the clone that is most prevalent within that cluster. All the CNA profiles are then compared to the scDNA CNA profile and reconstruction quality is measured by Pearson correlation coefficient.

#### Benchmarking CNV state prediction

We evaluate CNV state performance in each cluster by predicting the states (amplification or deletion) for all non-neutral bins of scDNA. We obtain the ground truth CNV states by running HMMs on the scDNA CNA profile. Echidna states are predicted by fitting HMMs on gene dosage *η* as well (for HMM detail see **HMM model for state prediction**). InferCNV states are determined by thresholding each cluster’s CNA profile using the cluster-specific standard deviation. Specifically, a bin is classified as amplification if it is one standard deviation or more above the mean, and as deletion if it is one standard deviation or more below the mean. Numbat states are derived directly from the clone-level pseudo-bulk profile output. For each bin, the maximum a posteriori (MAP) CNV states of all SNPs within the bin are collected, and majority voting is used to determine the bin’s final CNV state. Prediction performance in each cluster are measured by standard metrics, namely precision, recall, true positive rate, and false positive rate.

### 7.10 Application to Melanoma

We applied Echidna to a melanoma patient from the KEYNOTE-001 clinical trial previously described in [20]. To prepare the data for use in Echidna, we normalized both treatment timepoints together and performed Leiden clustering in order to get phenotypic states. Matched WGS data were processed by ichorCNA, as described in [20], and subsequently mapped to individual genes for use with Echidna.

For downstream analysis, we first computed DE genes by aggregating cells belonging to clones the Echidna classified as expanding or contracting. Specifically, a clone is labeled as expanding if the log2 fold change of the fraction of cells in that clone in the on timepoint divided by the fraction of cells in that clone in the pre timepoint is greater than 0.75, and contracting if the fold change is less than -0.75. (If a clone was only present in the pre-therapy time point, it is automatically labeled as shrinking, and vice versa if it is only present in the on time point).

For visualization **Fig. 5a, b**, we selected the top 100 significant DE genes in each condition, ranked by positive log fold change. The density of these genes along the genome was inputted into the Find Peaks function of Scipy. We defined a genomic hotspot to be a peak of DEGs over a region that contained a majority of amplified labels, as predicted by our HMM.

The significance of the GDX of all significant positive DEGs in each hotspot was compared against the full list of significant positive DEGs using a permutation test. The same permutation test was applied to study the significance of individual gene signatures in **Fig. S6**.

### 7.11 Analysis of Melanoma Spatial SlideSeq data

Spatial SlideSeq data previously published in [20] was loaded into Squidpy [61] and normalized using Scanpy. To determine the spatial localization of clones, we identified the top 20 differentially expressed genes ranked by log fold change in Clone 4 in the on-treatment timepoint. We colored the spatial plot by the z-score of the average expression across these genes, and visually identified the region of the plot that was enriched for the signature. We then computed the Moran’s I score for spatial autocorrelation as implemented by [61], on the set of highly varying genes returned by Echidna from the scRNA-seq. Genes from among those with the highest Moran’s I score were manually selected based on biological interest and range of GDX scores, and their z-scored expression was visualized in the spatial context.

### 7.12 Implementation and usage

The model is implemented with the PyTorch based statistical programming framework Pyro. The package follows the schematic of related tools for single-cell analyses, namely the `scanpy` package [66] and `decipher` [44]. The package manages the user’s analysis workflow through an `AnnData` object, systematically storing intermediate data and metadata as functions from the `.tl` submodule are called. Plotting functions in the `.pl` submodule automatically retrieve stored data from the `AnnData` object’s `.uns`, `.var`, and `.obs` attributes, simplifying data management for the user.

Typical usage begins by importing the package with `import echidna as ec`. The user has the option to toggle between single timepoint mode(if samples are collected at the same time point) and multi-timepoint mode(if samples are collected at different timepoints). To ensure a balanced representation of all time points, we downsample cells from each time point to the minimum number of cells across time points for multi-timepoint mode. The user trains the model by invoking `ec.tl.echidna train(adata, w df)`, which requires an `AnnData` object `adata` containing scRNA data and a pandas DataFrame `w df` with copy numbers for each time point, indexed by gene names. With a trained model, the user calculates learned clones by calling `ec.tl.echidna clones(adata)` and can then estimate growth status with `ec.tl.echidna status`. For investigating copy number aberrations across the genome, `ec.tl.echi cnv(adata, genome)` provides a comprehensive solution. This function accepts a pandas DataFrame `genome` with columns ‘chrom’, ‘txStart’, ‘txEnd’, ‘geneName’, and assigns amplification, neutral, or deletion status to each gene within a learned clone. While a default genome is provided, users are encouraged to use a genome most appropriate for their specific dataset. The results can be visualized using `ec.pl.plot cnv(adata)`. The package enables the calculation and visualization of gene dosage effects through `ec.tl.gene dosage effect(adata)` and `ec.pl.plot gene dosage(adata)`, respectively. For a comprehensive understanding of the package’s capabilities, users are directed to the example notebooks, which offer detailed, step-by-step walkthroughs.

## 8 Software and Code availability

The Python implementation of Echidna is available at https://github.com/azizilab/echidna. The notebooks to reproduce figures are available at https://github.com/azizilab/echidna reproducibility

